# Sea-ice microbial community succession and the potential role of parasitoids in the maintenance of diversity during the spring bloom

**DOI:** 10.1101/2025.05.18.654361

**Authors:** Kyle B. Dilliplaine, Laura M. Whitmore, Ana Aguilar-Islas, Channing Bolt, Kenneth Dumack, Meibing Jin, Mette Kaufman, Marc Oggier, Gwenn M. M. Hennon

## Abstract

Sea ice is a crucial, yet declining, habitat in high latitude ecosystems. Here we present a high-temporal resolution amplicon sequence data set collected during the spring ice-algal bloom near Utqiaġvik, Alaska in 2021 to study sea-ice microbial dynamics. The ice-algal bloom peaked on May 8^th^, reaching 46.6 mg chlorophyll *a* m^-2^, and thereafter became limited by nitrate availability. A massive bloom of the oil-degrading bacterium, *Oleispira* (>80% relative abundance), coincided with the algal bloom raising questions about hydrocarbon exposure. The sea-ice algal bloom was dominated by diatoms, particularly *Nitzschia spp*., and transitioned into a flagellate dominated post-bloom community which aligned with melt-associated changes to the physicochemical environment. We explored the relationship between putative parasitoids, Chytridiomycetes, Thecofilosea (Cercozoa), Oomycetes, Syndiniales (Dinoflagellata), and Labyrinthulomycetes (Bigyra), and potential microalgal hosts. Chytrids peaked periodically suggesting synchronized infections and *Cryothecomonas* (Thecofilosea) was observed parasitizing *Nitzschia spp*. for the first time in Arctic sea ice. Co-occurrence analysis suggested that diatoms, especially *Nitzschia*, were the primary hosts of Pacific-Arctic parasitoids, and that top-down parasitoid control may dramatically alter community composition over short timescales, such as days. These results provide important insights into the drivers of spring bloom timing and maintenance of microalgal diversity in sea ice.

## 1. Introduction

Climate change is rapidly impacting the Arctic sea-ice system by shifting the timing of freezeup and melt (Walsh et al., 2022), precipitation patterns (Bintanja and Selten, 2014; Webster et al., 2014), and ice extent and thickness (Babb et al., 2022; Stroeve and Notz, 2018). Sea-ice associated (sympagic) microalgae contribute substantially to Arctic primary production(Gosselin et al., 1997; Mortenson et al., 2020). Ice-algal production is primarily controlled by the availability of light, which is determined by ice and snow thickness (Mcdonald et al., 2015; Veyssière et al., 2022), and inorganic nutrients supplied via underlying water (Cota et al., 1987; Cota and Smith, 1991; Dalman et al., 2019). Sympagic communities have been observed to shift toward heterotrophic protists and dinoflagellates during the spring to summer transition (Rózańska et al., 2009; von Quillfeldt et al., 2003). Irradiance levels likely contribute to species-specific succession and community composition due to differing photophysiology, e.g., centric diatoms (such as *Thalassiosira* spp.), dinoflagellates, and certain pennate diatoms (such as *Navicula* spp.), increase with greater light availability while pennate diatoms like *Nitzschia frigida* decline (Croteau et al., 2022; Duncan et al., 2024). Nutrient availability can also influence microalgal community structure based on the resource utilization traits and stoichiometric requirements of different algal taxa (Litchman et al., 2015a, 2015b). Nitrogen is generally the limiting nutrient throughout most of the Arctic (Pineault et al., 2013; Rózańska et al., 2009), though silicic acid, which is required by diatoms, has been found to be limiting regionally (Giesbrecht and Varela, 2021; Gosselin and Legendre, 1990; Smith et al., 1990). Sympagic algae are positioned in the upper ocean, allowing them to take advantage of light early in the spring season, thus providing an early source of nutrition to the broader Arctic food web (Kohlbach et al., 2017). Sea ice melt processes release dense algal material which may be grazed by plankton (Juul-Pedersen et al., 2008), sink rapidly to the seafloor as aggregates (Ambrose et al., 2005; Boetius et al., 2013), or seed a successive phytoplankton bloom (Garrison et al., 1987; Lizotte, 2001; Yan et al., 2020). While much is known about bottom-up environmental control of sympagic microbial communities, top-down biological pressures may also affect community composition and are less well-understood.

The study of parasites in sea ice has been limited to small and focused investigations. While fungi such as chytrids (Hassett and Gradinger, 2016), and a newly discovered thraustochytrid (Labyrinthulomycetes), infect sea-ice algae (Hassett, 2020), Arctic Cryomonadida (Cercozoa, Thecofilosea), have thus far only been observed as phagotrophic grazers (Thaler and Lovejoy, 2012), though the commonly reported genus, *Cryothecomonas,* is currently the only known cryomonad genus to exhibit endoparasitic behavior (Drebes et al., 1996; Kühn et al., 2000). In contrast to parasites, parasitoids ultimately kill their host after infection (Skovgaard, 2014).

Parasitoids are capable of terminating phytoplankton blooms and altering species succession (Chambouvet et al., 2008; Tillmann et al., 1999). Despite their potential ecological importance, marine parasitoids are understudied due to ambiguous morphologies that make identification difficult (Käse et al., 2021), though molecular surveys suggest high undescribed diversity (B. T. Hassett et al., 2019; Thaler and Lovejoy, 2012). Identifying potential parasitoid-host interactions is required to better understand bloom dynamics and biogeochemical cycles in sea ice where parasitoid infections of microalgae have the potential to modulate the transfer of carbon between trophic levels (Amundsen et al., 2009).

Despite the importance of sympagic microbes to the Arctic ecosystem, basic information regarding microbial community structure, dynamics, and microalgal bloom phenology is spatiotemporally limited along the coastal US Arctic. We conducted a high-resolution time series (April 21 to June 11, 2021, sampled every few days) to determine the timing, magnitude, and yield-limiting factor of the ice-algal bloom at Utqiaġvik, Alaska, formerly known as Barrow. We examined the prokaryotic and unicellular eukaryotic community composition and dynamics over the spring microalgal bloom using 16S and 18S rRNA amplicon sequencing, and leveraged these data to identify putative parasitoid-host relationships within the sympagic community.

## 2. Methods

### 2.1. Study site and ice sampling

Field collections were conducted near Utqiaġvik, Alaska (71.375 N, 156.537 W) between April 21 and June 11, 2021 (Day of Year (DOY) 111 — 162), on a large level pan of landfast sea ice, measuring ∼11 km^2^ (Fig. S1). Additional site information and detailed methodology is provided in Appendix 1. Sea ice cores were collected from a ∼900 m^2^ area, with each sampling event constrained to 1 m^2^. Ice core samples were collected approximately three times per week and measurements of temperature, salinity, density, nutrients, chlorophyll *a* (chl *a*), and samples for DNA were collected (seeAppendix 1 for more details). Cores were retrieved using a 9-cm inner diameter Kovacs ice corer. Incident and under-ice photosynthetically active radiation (PAR; 400-700 nm) photon flux densities were measured adjacent to the sampling area using a LI-COR LI-1400 data logger with planar (incident; 2π) and spherical (under-ice; 4π) quantum sensors (Lincoln, NE, USA), respectively, and a 4π:2π ratio was used to determine “pseudotransmissivity” (see Appendix 1 for more details). Biological samples were obtained from the bottom 10 cm of a single ice core by quickly removing the section with a cleaned handsaw. The section was sealed in a clean plastic bag and stored in a cooler in the dark until transport back to the lab. Samples were processed by adding 100 ml of 0.2 µm filtered seawater cm^-1^ of ice to prevent osmotic shock (Garrison and Buck, 1986) and melted at 4 °C in the dark overnight. Once entirely melted, 1-2 aliquots (12-171 mL) of each sample were vacuum filtered (≤ 5 psi) for chl *a* using 25 mm glass fiber filters (Whatman GF/F). All duplicated chlorophyll subsamples were averaged and reported as integrated areal chl *a* concentrations. The remaining sample volume was processed for DNA by filtering the sample onto a 0.2 µm Sterivex filter (Millipore Sigma) using a peristaltic pump and stored at -80 °C until further processing.

### 2.2. DNA extraction, sequencing, and bioinformatic processing

DNA was extracted using the NucleoMag DNA/RNA Water kit (Macherey Nagel, Düren, GE). The V4 region of both the 16S and 18S rRNA genes were amplified for each sample using the revised Earth Microbiome Project primers (515FB and 806RB; Caporaso et al. 2012; Apprill et al. 2015; Parada et al. 2016) and TAReuk454FWD1 and TAReukREV3 (Stoeck et al., 2010), respectively. Primers were modified with TaggiMatrix indices to enable pooling of samples prior to adaptation with Illumina sequencing adapters (Glenn et al., 2019). Samples were sequenced on an Illumina MiSeq with a 2 x 300 paired-end kit at the University of Alaska Fairbanks’ Genomics Core Laboratory. The QIIME2 (version 2023.5) workflow was used to process reads before further analysis (Caporaso et al., 2010); pre-processing and detailed information can be found in Appendix 1. Prokaryotic amplicon sequence variants (ASVs) were classified using a weighted classifier (515f-806r-average-classifier.qza) trained on the Silva 138.1 database (Bokulich et al., 2018; Kaehler et al., 2019; Quast et al., 2012; Robeson et al., 2021). A classifier was trained on the PR2 database (version 5.0.0) for eukaryotic ASV classification (Guillou et al., 2012). ASVs were identified to the lowest taxonomic level possible and ASVs of the same taxonomic identity were numbered sequentially (Table S1). Taxonomic corrections were made to the eukaryotic classifications due to classifier uncertainty and incomplete taxonomic representation in the PR2 database (Table S1). To provide the most up-to-date taxonomic assignment, each ASV involved with a significant parasitoid correlation was manually checked via NCBI BLAST alignment. Additional details regarding sequence processing can be found in Appendix 1.

### 2.3 Bioinformatic analysis

Community structuring across our time-series was investigated according to Walker et al. (2023) for rarefied prokaryotic and eukaryotic ASV data. Community structure was visualized using non-metric multidimensional scaling (nMDS) using Bray-Curtis dissimilarity matrices with Wisconsin double standardization transformed ASV tables. Outliers were identified using Mahalanobis distance using ‘rstatix’ (Kassambara, 2023); outliers were retained on the nMDS plot but were excluded from cluster polygons. Cluster association indices were calculated for each ASV to identify which were significantly associated with each of the clusters. The ‘indicspecies’ package was used to calculate the point biserial correlation coefficient on rarefied data (Cáceres and Legendre, 2009), and corrected for unequal sample distribution between clusters (Tichy and Chytry, 2006). Only taxa with *p* < 0.001, correlation coefficients >0.5, and contributed at least 0.5% of all sequences (16S or 18S) were retained.

Correlations between putative parasitoids and phototrophic hosts were investigated using Spearman’s rank correlation of select ASVs using the ‘psych’ (Revelle, 2023; v.2.3.9) package in R with false discovery rate corrections for *p* values (Benjamini et al., 2009). Unrarefied relative abundance data were first centered log-ratio transformed using the ‘SpiecEasi’ package (v.1.1.3; Kurtz et al., 2017), after removal of rare taxa (occurring as <0.1% in fewer than 50% of samples). Parasitoids were limited to the classes Chytridiomycetes, Thecofilosea (Cercozoa), Labyrinthulomycetes (Bigyra), Oomycetes and Syndiniales (Dinoflagellata) while hosts were limited to the dominant phototrophs, i.e., diatoms (Bacillariophyceae, Mediophyceae, and Coscinodiscophyceae) and dinoflagellates (Dinophyceae). To assess whether parasitoids are associated with microalgal α-diversity (limited to the taxa defined above), we regressed the per-sample Inverse Simpson index on the summed relative abundance of all parasitoid classes. Correlation matrices were produced at the ASV level, retaining only those with at least a moderate correlation strength, i.e. |rho| ≥ 0.5; *p* < 0.05). We sampled again in April of 2022 and used microscopy to validate potential correlations and quantify *Cryothecomonas* infection prevalence (see Appendix 1for more details).

## 3. Results

Snow depth ranged from 0 to 6 cm during the study period (Figs. 1c & S2b). Direct measurements of under-ice photosynthetically active radiation (PAR) ranged from 2.34 – 77.50 µmol photons m^-2^ s^-1^ (Fig. 1c) which was 0.25 – 11.13% of incident PAR (Fig. S2). The air temperature began warming on DOY 133 and reached above 0 °C on DOY 141 (Fig. S3a). The temperature measured at 2.5 cm from the bottom of the ice warmed above -1.8 °C by DOY 134, indicating the initiation of bottom ice melt between DOY 132 and 134 (Fig. S3d; ice growth < DOY 134). Ice thickness was relatively uniform with a thickness of 109 ± 2 cm (mean ± SD) until melt onset, declining thereafter to a minimum thickness of 95 cm (Fig. S2c). The brine salinity from the bottom 10 cm decreased substantially between DOYs 132 and 137 (Fig. S4c). The peak chlorophyll biomass occurred on DOY 128 (May 8^th^), reaching a maximum of 46.6 mg chl *a* m^-2^ (Fig. 1). The calculated accumulation and loss rates were 2.64 and -4.57 mg chl *a* m^2^d ^-1^, respectively. Areal chl *a* concentration declined below 1 mg m^-2^ on DOY 144 (May 24^th^); bloom phases are therefore defined as “bloom” < DOY 144 and “post-bloom” ≥ DOY 144 (Fig. 1a). The average proportion of chlorophyll degradation product (phaeophytin) increased from 31% to 45% after the onset of bottom ice melt. Seawater nutrient concentrations peaked on the date of bloom termination (DOY 144; Fig. 1b). Bulk nutrient concentrations in the bottom 10 cm of ice were significantly correlated with chl *a*; nitrate (adj. r^2^=0.45, *p* =0.0014), nitrite (adj. r^2^=0.17, *p* =0.0447), ammonium (adj. r^2^=0.36, *p* =0.0038), phosphate (adj. r^2^=0.63, *p* <0.0001) and silicate (adj. r^2^=0.41, *p* <0.0019; Fig. S5).

**Figure 1.**
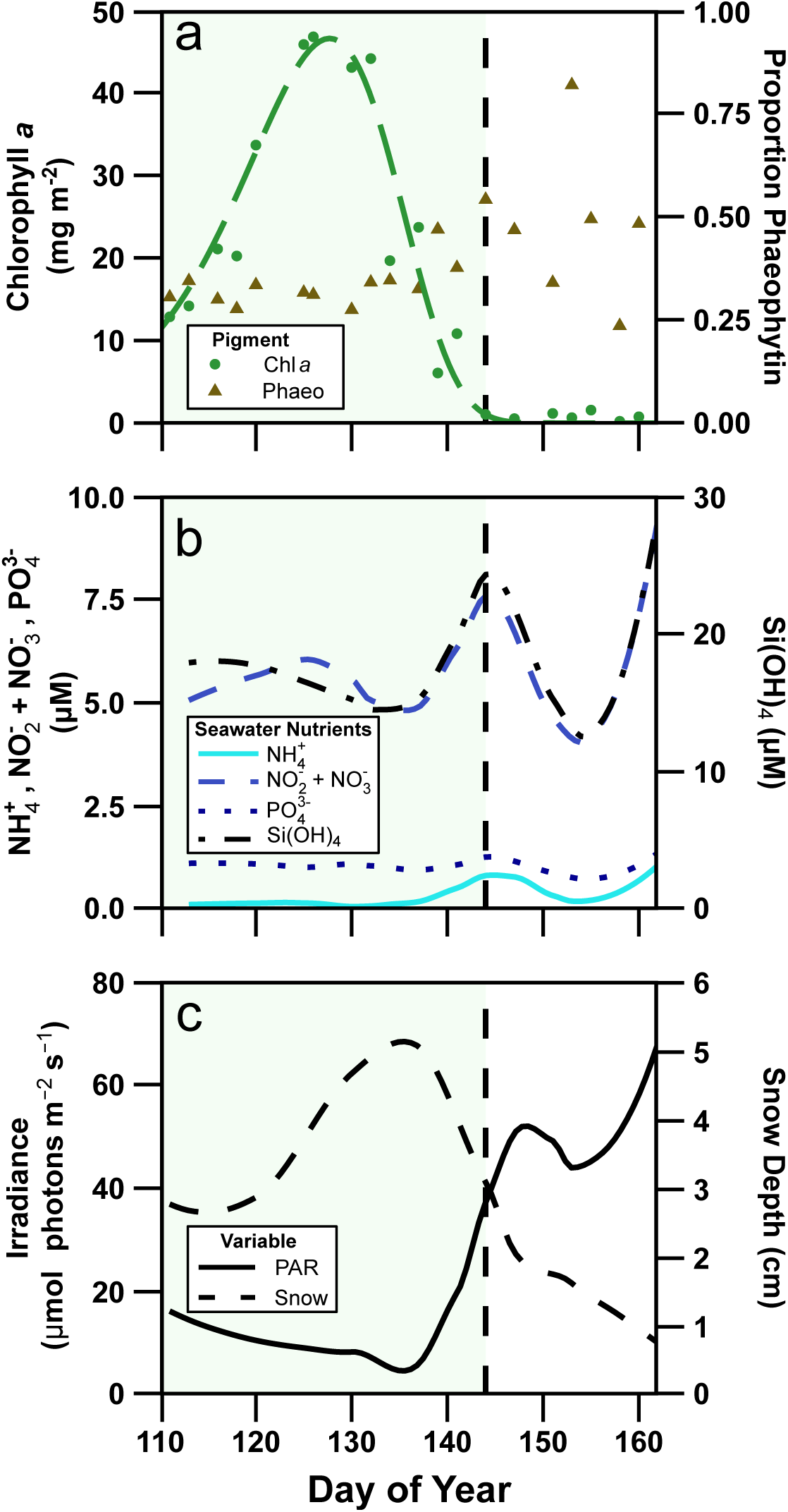
Development of biologically relevant measures over time. Temporal evolution of chlorophyll *a* and phaeophytin proportion (a), seawater concentrations of ammonium (NH_4_^+^), nitrite + nitrate (NO_2_^-^+ NO_3_^-^), phosphate (PO_4_^3-^) and silicate (Si(OH)_4_), (b), and snow thickness and under-ice PAR as measured by a spherical quantum sensor (4π; c). Green shading indicates the ice-algal bloom and the vertical dashed line indicates the bloom termination (<1 mg chlorophyll *a* m^-2^).

### 3.1.1. Sympagic community composition patterns

The total number of prokaryotic ASVs identified from the unrarefied data was 469, representing 124 families from 33 classes of bacteria and archaea. A relative abundance of approximately 48% of the sequence reads belonged to just two ASVs identified to the Genus level, *Oleispira* sp. and *Paraglaciecola* sp., 30% and 17.5%, respectively. *Polaribacter* was most abundant early in the time-series before transitioning to a bloom of *Oleispira* that approximately follows the microalgal bloom (Fig. 2). *Tenacibaculum* and *Colwellia* genera were the most abundant taxa in the post-bloom period (Fig. 2). The combined relative abundance of all four *Oleispira* ASVs reached a peak of 84.4% on DOY 120 and was above 5% during a 35-d period (n=15); eight of those samples were above 50% (Fig. S6a). A positive correlation was found between *Oleispira* relative abundance and the areal chl *a* concentration (adj. r^2^ = 0.68, *p* = <0.001; Fig. S6b).

**Figure 2.**
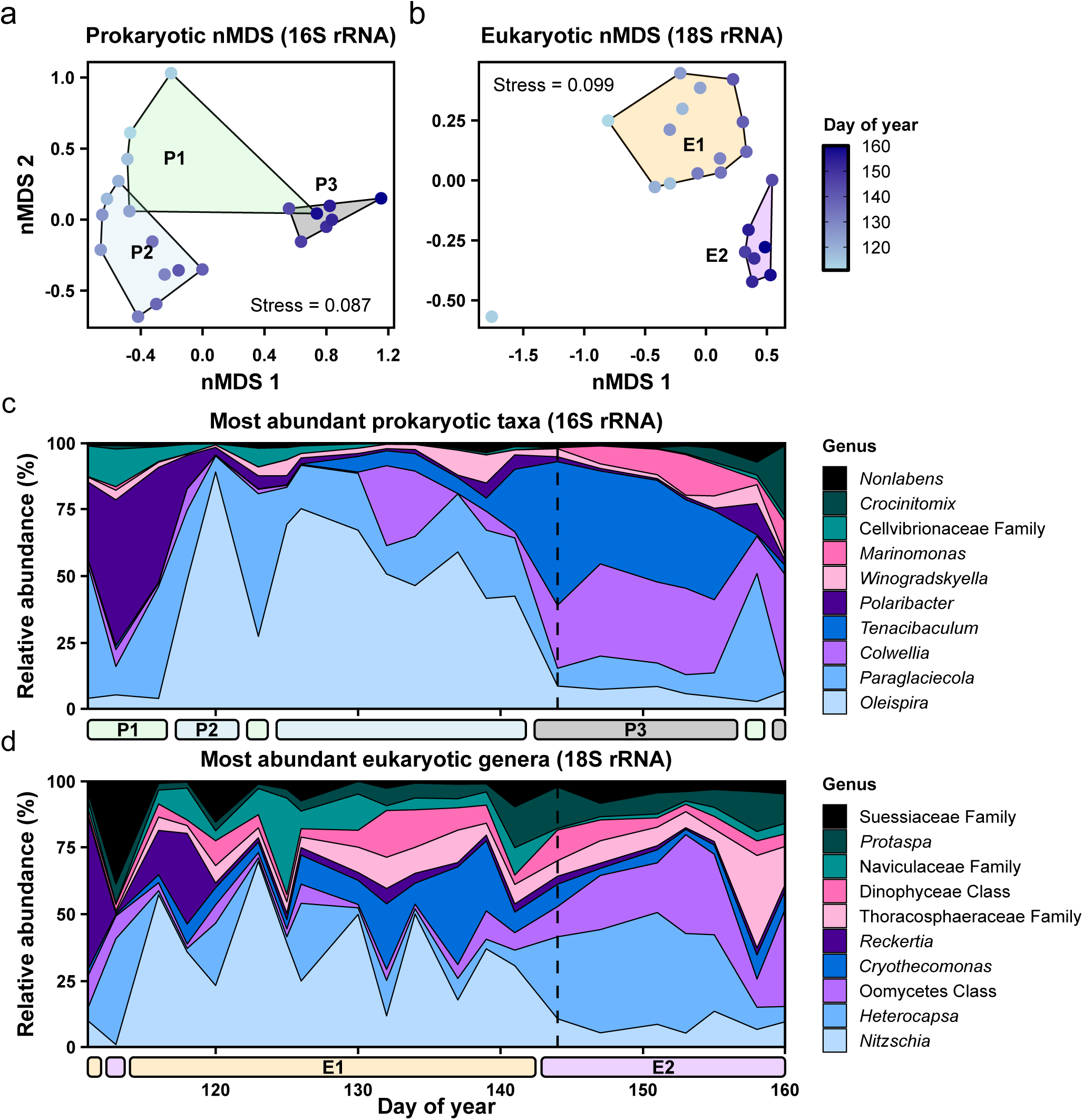
Sympagic microbial community structure and the most abundant genera over time. Non-metric multidimensional scaling (nMDS) ordination using Bray-Curtis dissimilarities (a & b). Sample groupings were determined by hierarchical clustering and labeled by their relation to the stage of the bloom. For prokaryotes (a), early- (P1), mid- (P2), and post-bloom (P3) clusters are observed. For Eukaryotes (b), bloom (E1) and post-bloom (E2) clusters were distinguished. Relative read contribution of the top 10 most abundant genera of prokaryotes (c) and unicellular eukaryotes (d). Vertical dashed line indicates bloom termination (<1 mg chlorophyll *a* m^-2^).

Three prokaryotic community clusters were identified as unique using hierarchical clustering analysis (adonis2: F = 8.59, *p* < 0.001), and was supported by a marginally insignificant within-cluster dispersion (betadisper: *p* = 0.075). The average silhouette width of 0.47 supports the separation of these clusters, referred to as clusters P1, P2, and P3 (Figs. 2a & S7a). Prokaryotic taxa identified by indicator taxa analysis that were significantly associated with the clusters are used as representatives for describing the major successional patterns and associated metabolisms (Table S3). *Polaribacter* was the dominant genus associated during the early-bloom (P1), preceding a bloom of *Oleispira* coincident with the microalgal bloom (P2), before transitioning to a more even post-bloom (P3) community represented by associated members of *Colwellia, Marinomonas,* and *Tenacibaculum*.

The total number of unicellular eukaryotic ASVs identified from the unrarefied data was 751, representing 14 divisions. The relative abundance of the top five most abundant classes contributed 75.4% of all sequences reads: Bacillariophyceae (26.5%), Dinophyceae (23.4%), Thecofilosea (13.8%), Oomycetes (6.8%), and the Imbricatea (4.9%). A total of 119 diatom ASVs contributed to 28% of the relative read abundance. The diatom genus *Nitzschia* contributed most to the bloom-phase, before transitioning into a post-bloom phase with a large contribution by the dinoflagellate genus *Heterocapsa* (Fig. 2d). Hierarchical clustering analysis identified two unique eukaryotic clusters (adonis2: F = 8.72, *p* < 0.001). The dendrogram indicated a substructure within these two major clusters (Fig. S7b). Within-cluster dispersion was not significant (betadisper: *p* = 0.292), and the average silhouette width of 0.27 is generally considered weak. We designate these two clusters as E1 (bloom) and E2 (post-bloom; Fig. 1c). One outlier (DOY = 113) was identified using Mahalanobis distance and retained in analyses, although excluded from the nMDS polygon (Fig. 2b). Eukaryotic association indices revealed taxa significantly associated with each of the two clusters (Table S4). Four diatom, and one flagellate, ASVs were associated with cluster E1, while seven flagellate ASVs were associated with cluster E2.

### 3.1.2. Parasitoid-host associations

The putative parasitoids investigated in this study appear to follow successional co-exclusion during the sympagic algal bloom, seen by their relative abundance (Fig. 3). The Chytridiomycetes followed a rhythmic pattern of peaks and troughs, roughly following a single cycle approximately every seven days during the bloom period. The Thecofilosea had genera dependent development during the time series, *Cryothecomonas* peaked during the late-bloom period while *Protaspa* became most prevalent after *Cryothecomonas* declined. The unclassified Cryomonadida (Cercozoa, Thecofilosea), Labyrinthulomycetes and Syndiniales remained relatively low throughout the timeseries, while the relative abundance of Oomycetes dramatically increased during the post-bloom period. Inverse Simpson index showed a weak (*r*^2^ = 0.07), insignificant (*p* = 0.2314) positive association with summed parasitoid counts (β = 0.15 ± 0.12 SE; Fig. S8). Directed correlations, i.e., parasitoid versus microalgal pairings, were investigated at the ASV level (Fig. 4). Of the 91 significant correlations, 75 were with ASVs of the Thecofilosea (62 of which were with diatoms). Using the combined positive and negative correlations as an indicator of host range, the thecofilosean parasitoids had the largest potential host range while the labyrinthulids were the most limited (Fig. 4). Special attention was given to the correlated ASV pairing of Lobulomycetales Order_1 with *Nitzschia* ASVs (Fig. 5). Subsequent microscope observations support at least some of the correlations that were found; to the best of our knowledge,we provide first-time evidence of internal parasitization of Arctic diatoms by *Cryothecomonas* (Fig. 6). In 2022, 5.1% ± 0.98% (mean ± SD) of *N. arctica* cells were found to be infected by *Cryothecomonas*.

**Figure 3.**
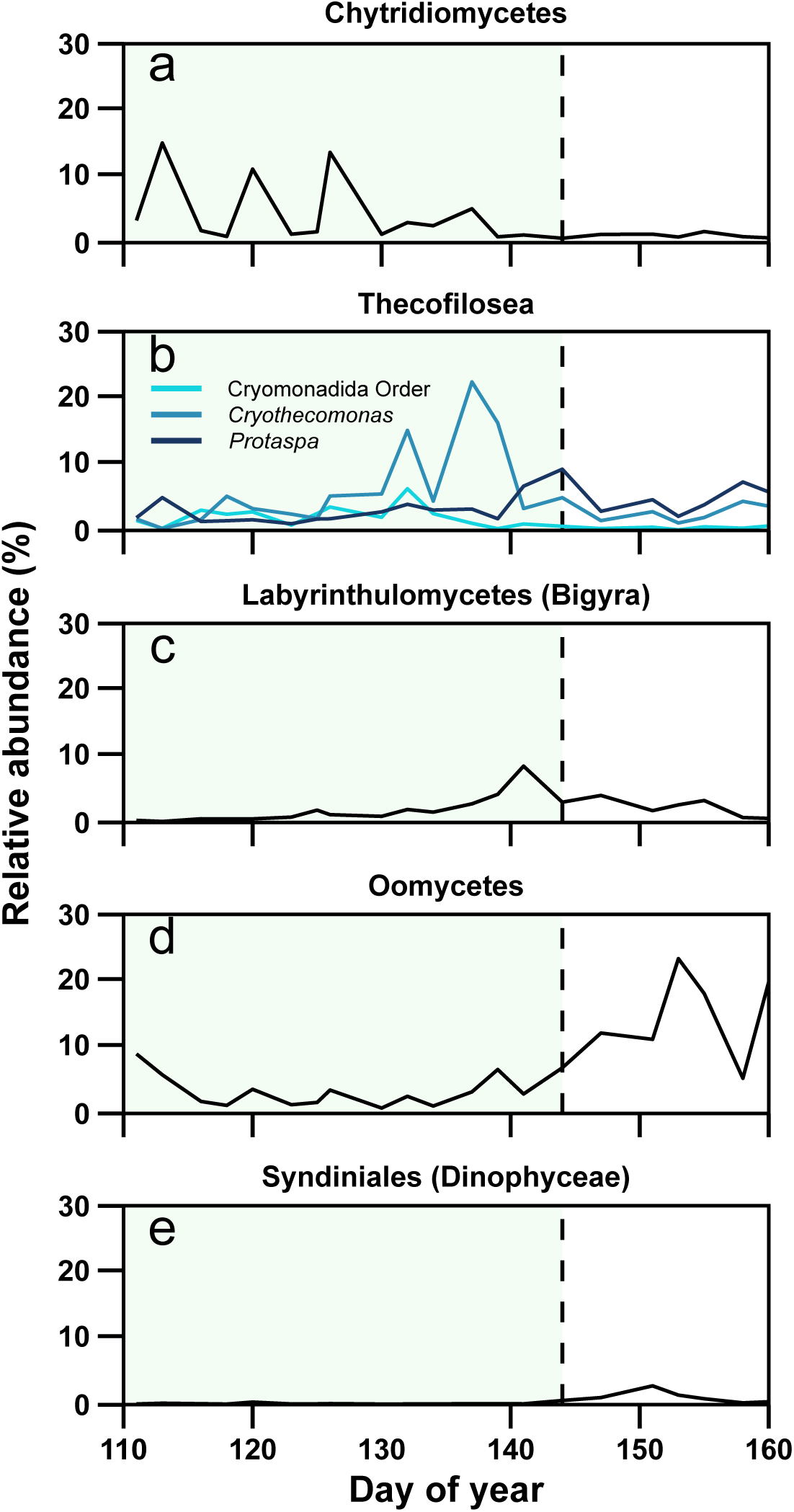
Relative abundance of putative parasitoid groups over time. (a) Chytridiomycota, class Chytridiomycetes, (b) Cercozoa, class Thecofilosea, (c) Bigyra, order Labyrinthulomycetes, (d) Oomycetes, (e) Dinoflagellata order Syndiniales. Green shading indicates the ice-algal bloom and the vertical dashed line indicates the bloom termination (1 mg chlorophyll *a* m^-2^).

**Figure 4.**
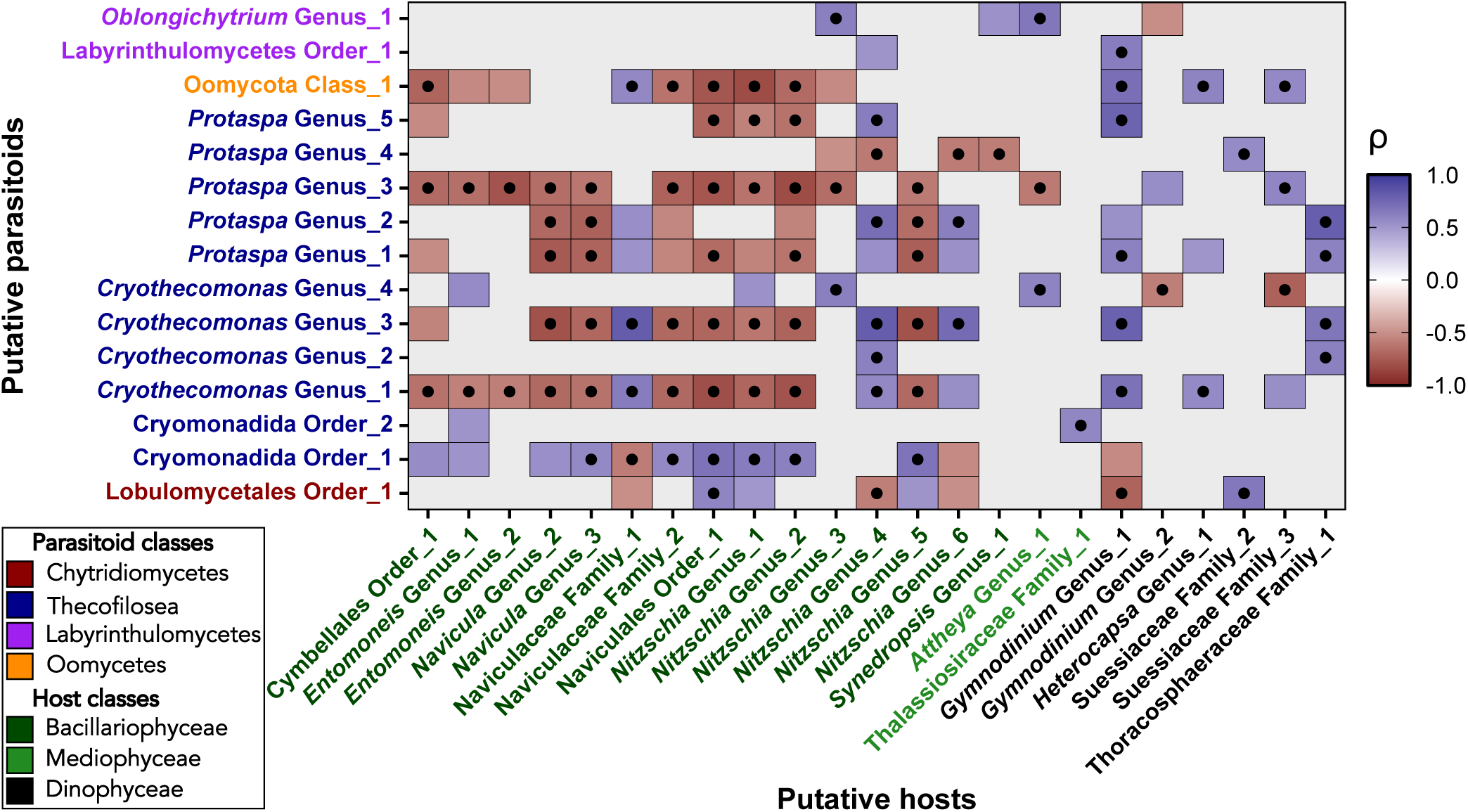
Correlation of putative parasitoids with putative hosts. Directed spearman correlation coefficient grid of amplicon sequence variants (ASVs) containing correlations between putative parasitoids and putative hosts with |rho| ≥ 0.5. Black dots indicate significant correlation (adj. *p* < 0.05).

**Figure 5.**
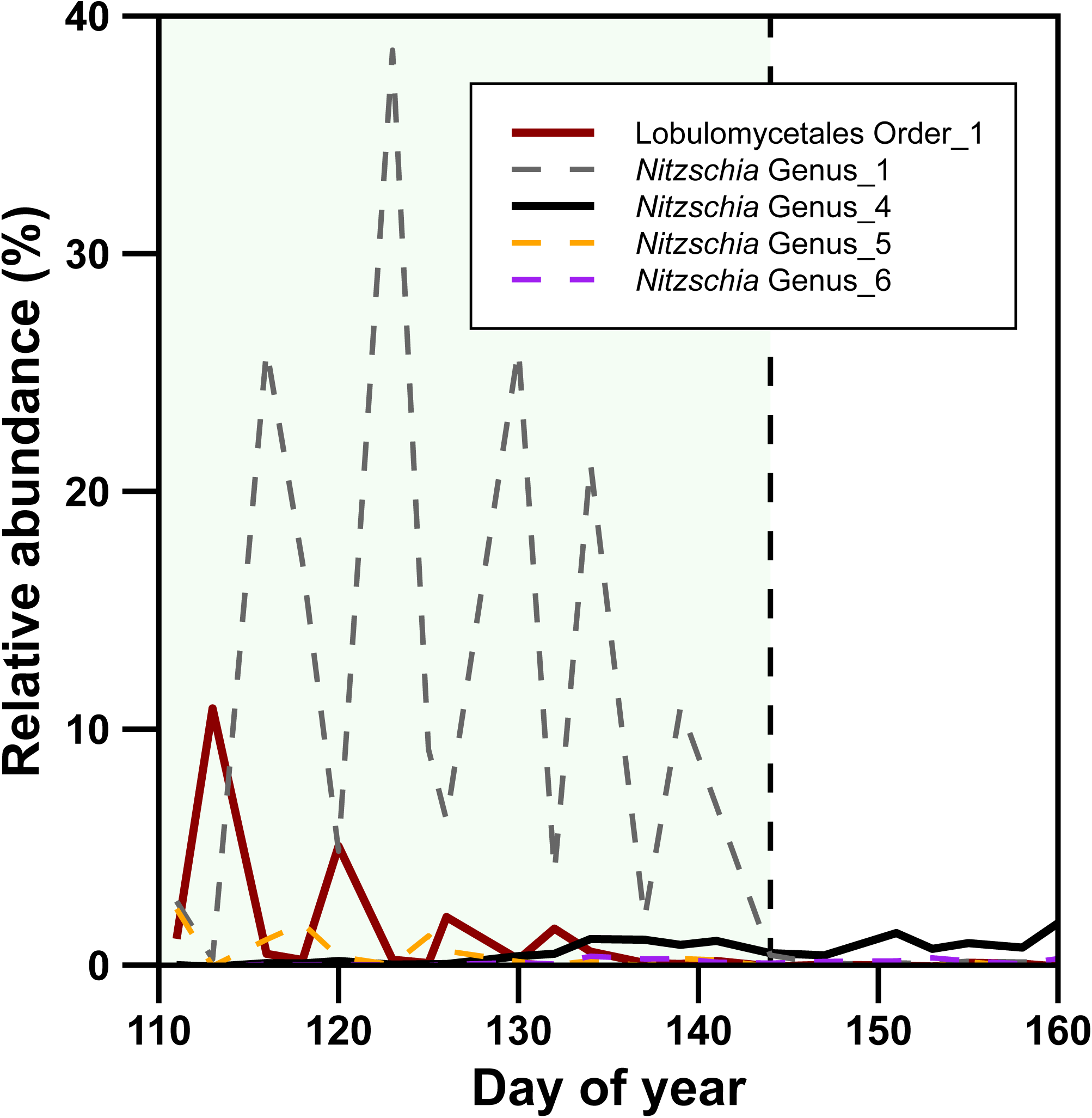
Relative abundance over time of a putative parasitoid (red line, Chytridiomycota) and *Nitzschia* hosts (black, grey, orange and purple lines). Solid lines indicate a significant correlation (p < 0.05); all *Nitzschia* ASVs presented here had a correlation coefficient ≥ |0.5|. Green shading indicates the ice-algal bloom and the vertical dashed line indicates the bloom termination (1 mg chlorophyll *a* m^2^).

**Figure 6.**
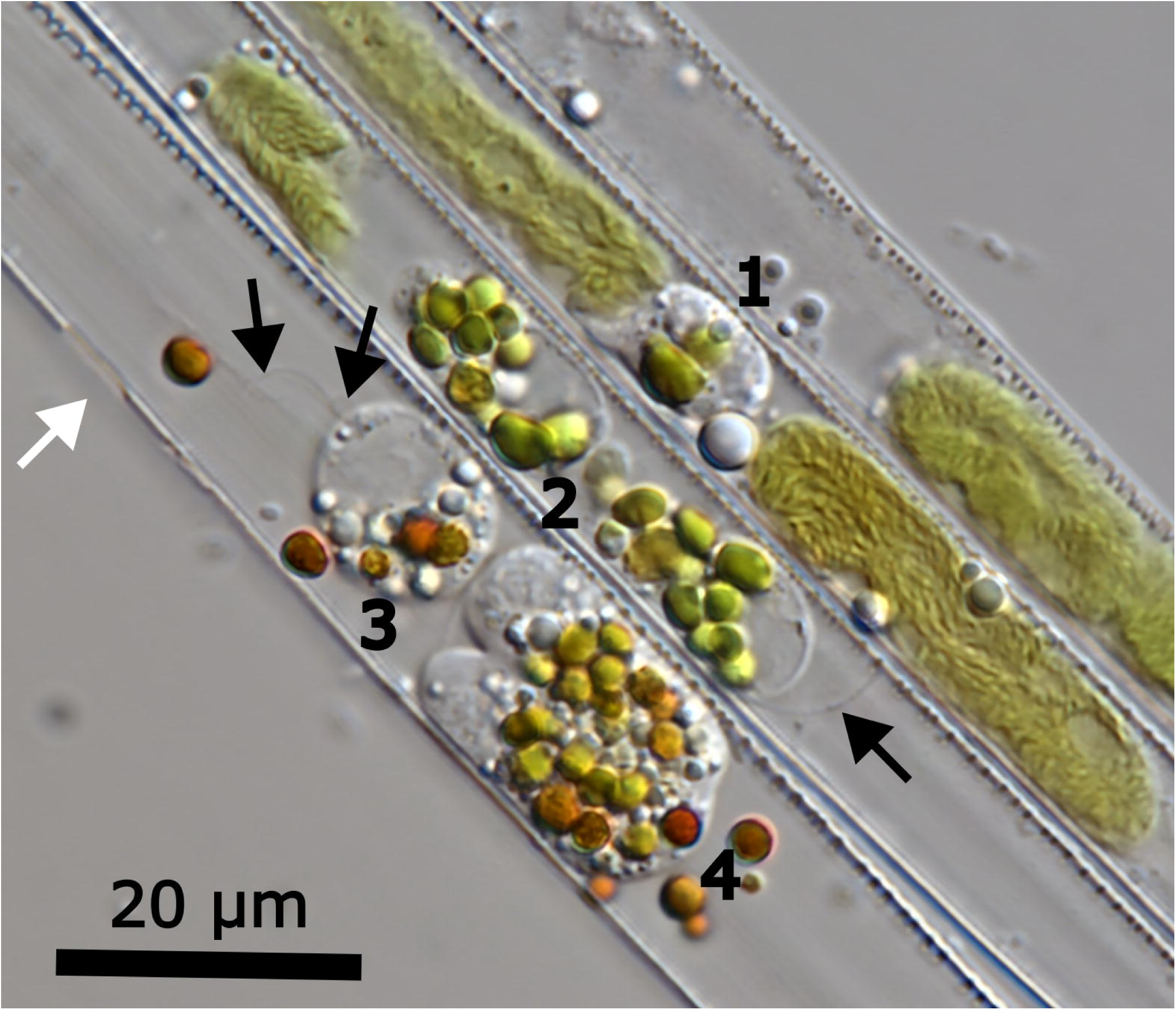
A small chain of Nitzschia arctica infected by *Cryothecomonas sp*. at different stages of infection. White arrow points to a compromise in the frustule that may be an entry point for the flagellate and black arrows point to flagella. 1) A single flagellate beginning to phagocytize the plastid of a newly infected diatom. 2) Two small daughter cells with digestive vacuoles and fresh, still green, plastid material. Note the partially consumed plastid in the top left of the diatom frustule. 3) A mature flagellate with brown digestive vacuoles and surrounded by several defecated brown fecal bodies. 4) A large flagellate with many brown digestive vacuoles (the missing diatom plastids) with a cleavage furrow developing at the top left indicating the initiation of longitudinal mitotic division.

## 4. Discussion

### 4.1. Physicochemical environment and bloom dynamics

The maximum ice thickness of undeformed sea ice at Utqiaġvik was approximately 0.5 m less than measurements from the early 2000’s (Jin et al., 2006; Mahoney et al., 2007; Lee et al., 2008; Manes and Gradinger, 2009; Fig. S2). The maximum annual landfast ice thickness near this location has thinned at a rate of 10 cm decade^-1^, over twice the rate of several locations in the Canadian Arctic Archipelago (Eicken et al., 2012; Howell et al., 2016; Osborne et al., 2018). Snow depth in 2021 was also thinner than previous years, which often reached ∼20 cm (or greater in drifts; Herzfeld et al., 2006; Jin et al., 2006; Manes and Gradinger, 2009). Thinner snow and ice layers provided greater light transmittance than in past decades (Figs. 1c & S2). Pseudotransmitted irradiances remained below photoinhibitory thresholds until the post-bloom period (∼50 µmol photons m^-2^ s^-1^; Arrigo et al., 2010; Juhl and Krembs, 2010; Croteau et al., 2022; Dilliplaine and Hennon, 2023; Fig. S2e), though, excluding chlorophyll from attenuation calculations suggests photoinhibition may have occurred at the top of the dense ice algal layer (Fig. S2h). An overall brighter under-ice light field is likely to become more common as ice thins and less snow falls, which may influence community composition or the vertical distribution of species as they balance light and nutrient requirements and preferences (Aumack et al., 2014; Croteau et al., 2022; Lannuzel et al., 2020).

The underlying seawater nutrient concentration and flux are generally considered important factors controlling the magnitude of ice algal blooms (Cota et al., 1991; Dalman et al., 2019; Gradinger, 2009; Rózańska et al., 2009). During our study, seawater nutrient concentrations (Fig. 1b) and nutrient ratios (Fig. S9) were similar to previous studies in the region during the spring ice-algal bloom (Lee et al., 2008; Manes and Gradinger, 2009). Nutrients remained replete in the underlying seawater, with resupply likely driven by unmeasured physical processes such as turbulence at the ice-water interface. We excluded the use of sea-ice brine nutrient data because it appeared that cell lysis during melting released intracellular nutrient pools, evident by the strong positive correlation of each macronutrient analyte with chl *a* (Fig. S5). Brine nutrient concentrations are expected to behave conservatively according to the equations of Cox and Weeks (1983), though Roukaerts et al. (2021) presented a biofilm-mediated hypothesis to explain the accumulation of nutrients with algae in the Southern Ocean. However, recent studies show that sea-ice nutrient-chlorophyll correlations likely reflect the release of intracellular nutrient pools during melting protocols (Mundy et al., 2025; Ahmed et al. in review). The positive silicate-chlorophyll relationship in our study suggests that our ∼24h direct melt method released contents of the silicon deposition vesicles. The observed nutrient ratios in underlying seawater were well below the expected 16:1 (N:P) from Redfield and greater than 1:1 (Si:N; Fig. S9), indicating nitrate resupply became the limiter to maximum biomass accumulation (Redfield et al., 1963) prior to the onset of melt. Using the empirical nitrate-chlorophyll relationship from Rózańska et al. (2009) the average seawater nitrate concentration (5.6 µmol L^-1^) predicts a maximum chl *a* concentration of 44.9 mg m^2^, closely matching our maximum observed concentration of 46.6 mg m^-2^ (Fig. 1a).

This study is the first to document the near-complete growth, decay, and fully realized magnitude of the sea-ice algal bloom at Utqiaġvik, AK, USA. The peak of the algal bloom occurred on DOY 128 (May 8^th^) similar to the peak date (DOY 121; May 1^st^) derived from regionally compiled data presented in Leu et al. (2015). The 2021 bloom magnitude exceeded all previously reported values near our study site, but was similar to collated data for landfast ice in Resolute Bay (Leu et al., 2015), and exceeds that of other landfast ice time-series such as the Green Edge ice camps in Baffin Bay (Massicotte et al., 2020; ∼30 mg m^-2^ 2015, ∼7 mg m^-2^ 2016). The highest prior measurement at this location was 36.2 mg chl *a* m□², measured in late May (Gradinger et al., 2009), while our peak was ∼25% greater than this isolated observation, and nearly 300% greater than the model estimate from compiled data (Leu et al., 2015). It remains unclear whether this elevated biomass reflects environmental change (e.g., thinning ice and snow, faster under-ice currents) or is due to higher temporal resolution. Sampling just one week before or after the actual peak would have underestimated biomass by 26-38%, respectively. If sea-ice algal biomass is increasing, it could lead to greater secondary production. Bloom termination and succession were tightly linked to the onset of sea ice melt, consistent with previous observations (Leu et al., 2015 and references therein; Oziel et al., 2019). Bottom-ice melt coincided with the increase of phaeophytin proportion (Fig. 1a), indicating the onset of unfavorable conditions such as freshening, nutrient depletion, grazing, or infection.

### 4.2. Sympagic community dynamics

We observed distinct patterns of succession in eukaryotic and prokaryotic communities associated with bloom stage (Fig. 2). The eukaryotic community gradually transitioned between the bloom (E1) and post-bloom (E2) phases, potentially driven by brine drainage and export of non-motile taxa. The E1 cluster was enriched in pennate diatoms and transitioned to flagellate dominance in E2 (Fig. 2; Table S4), a well-established progression in Arctic sea ice (Alou-Font et al., 2013; Rózańska et al., 2009; von Quillfeldt et al., 2003).

We observed three distinct prokaryotic community clusters, largely following the temporal evolution of the bloom: early- (P1), mid- (P2), and post-bloom (P3; Figs. 2a). Such changes are likely driven by shifts in dissolved organic matter availability and composition, as prokaryotic communities respond to the dynamic concentration and quality of organic matter that evolve over the course of microalgal bloom development(Zhang et al., 2018; Zhou et al., 2018). The P1 associated *Polaribacter*, and P3 associated *Tenacibaculum* and *Colwellia* ASVs, are heterotrophic bacteria known for their utilization of microalgal-derived dissolved organic matter (DOM; Landa et al., 2016; Underwood et al., 2019). *Colwellia* and *Tenacibaculum* are capable of degrading high molecular weight DOM; additionally, *Tenacibaculum* can utilize complex extracellular polymeric substances derived from diatom biofilms (Bohórquez et al., 2017; Underwood et al., 2019). *Marinomonas,* which was also associated with the P3 cluster, has the capacity to metabolize aromatics from terrestrial peat (Sipler et al., 2017). These results suggest *Polaribacter* uses labile carbohydrates, while P3 members degrade recalcitrant high-molecular-weight substrates remaining after labile material is consumed. *Paraglaciecola* and *Winogradsyella*, which have been previously observed associated with sea-ice algal aggregates (Rapp et al., 2018), were abundant throughout the time series and not associated with any of the clusters. This may be due to their ability to use a diverse array of carbohydrates (Schultz-Johansen et al., 2018; Sun et al., 2025), maintaining abundances as the quality and quantity of algal substrates change.

During the mid-bloom (P2), the prokaryotic community was unexpectedly dominated by the oil-degrading bacterium *Oleispira* (Fig. S6; Table S3). While many sympagic genera (e.g., *Colwellia, Polaribacter, Nonlabens, Paraglaciecola*) can degrade hydrocarbons, *Oleispira* is an obligate hydrocarbonoclastic bacterium that rapidly blooms after petroleum exposure (Brakstad et al., 2008; Lofthus et al., 2018; Netzer et al., 2018; Yakimov et al., 2003; Yang et al., 2016). Relative abundances >5% typically indicate hydrocarbon input (Krolicka et al., 2019), and although *Oleispira* exceeded 80% in some samples there was no visible contamination. This enrichment may reflect advection from a distant spill and subsequent entrainment into bottom ice via seawater flux and physical sieving (Spindler, 1994; Syvertsen, 1991).However, the close association of hydrocarbonoclastic bacteria, such as *Oleispira*, with phytoplankton (Thompson et al., 2020; this study), or decaying microalgae (Cono et al., 2022), and the correlation of abundance with chl *a*, warrants further evaluation of *Oleispira* as an indicator taxon of fossil hydrocarbon exposure, as these could be sources of biogenic hydrocarbons.

The eukaryotic cluster, E1, was associated with the diatom genera *Nitzschia* and *Navicula,* which are among the most abundant algae in Arctic sea ice (Table S4; Leu et al., 2015). An unidentified cryomonad (Cryomonadida Order_1) was also associated with E1; Cryomonadida are commonly reported in Arctic sea ice and are generally considered heterotrophic (Thaler and Lovejoy, 2012). Rhythmic fluctuations of *Nitzschia* relative abundance were seemingly unrelated to environmental drivers. These fluctuations may be explained by spatial heterogeneity, or parasitoid dynamics as discussed in section 4.3.1.

The E2 community was associated with heterotrophic flagellates and ciliates (Table S4), consistent with summer sea ice (Leu et al., 2015; Marquardt et al., 2023). Dinoflagellates included *Gymnodinium* and *Heterocapsa*with *Gymnodinium* spp. known to feed on small diatoms (Eddie et al., 2010) and are correlated with higher light conditions (Duncan et al., 2024). Additional E2 ASV associations included included a Pyramimonodaophyte, *Pyramimonas sp.,* thecofilosean *Protaspa*, Oomycetes and Cilliophora. The green alga, *Pyramimonas*, sometimes forms dense blooms in under-ice brackish water ponds associated with melt conditions (Gradinger, 1996; Massicotte et al., 2020). The heterotrophic flagellate *Protaspa* can feed on diatoms using pseudopodia (Howe et al., 2011; Schnepf and Kühn, 2000). The Oomycetes can be parasitic (Thines et al., 2015) or saprotrophic, thus their increased presence in the post-bloom community may represent an opportunity to feed on light and salinity stressed diatoms or on the post-bloom exopolymers that are preferentially retained during melt (Juhl et al., 2011; Meiners et al., 2008). The abundance of putative parasitoids both in bloom and post-bloom conditions was striking, leading us to further explore their potential role in shaping microalgal succession. While prokaryotic community composition appears largely shaped by bottom-up processes, with shifts in dissolved organic matter quality and quantity selecting for taxa with specialized metabolic capacities, eukaryotic community composition reflects both bottom-up drivers, e.g., nutrient and light availability and desalination processes, and top-down interactions such as those from parasitoids and grazers.

### 4.3. Sea-ice parasitoid dynamics and potential host range

To explore the relationship of parasitoid groups in shaping sea-ice microalgal communities, we identified putative parasitoid sequences from those taxonomic groups expected to be found in relatively high proportion within our samples: Chytridiomycetes (Fungi), Cryomonadida (Cercozoa, Thecofilosea), Oomycetes, Syndiniales (Dinophyceae) and Labyrinthulomycetes (Bigyra). Within the Cryomonadida, the relationship between *Protaspa* and *Cryothecomonas* are not fully resolved, so both of these groups were retained in this study despite parasitism confined only to the *Cryothecomonas*. Our time-series data revealed successional patterns through the microalgal bloom (Fig. 3). Environmental controls on parasitoid infections remain poorly understood, aside from a few specific cases. Infections by the cryomonad *Cryothecomonas aestivalis* appear inhibited below 4 °C (Catlett et al., 2023; Peacock et al., 2014). Our observations demonstrate that parasitoid groups periodically spike, suggesting that infections are occurring throughout the spring bloom at temperatures below -1 °C (Fig. 3). Brine salinity varies more widely in the spring (Fig. S4c) and likely plays a key role in regulating parasitoid pathogenicity in sea ice. Chytridiomycetes were abundant during the early bloom but declined with melt onset and may be stenohaline, while cryomonads and Oomycetes may be euryhaline and tolerant of fresher meltwater (Fig. 3). However, previous studies report both cryomonads and chytrids as more common in phytoplankton influenced by sea-ice melt waters (Kilias et al., 2020; Thaler and Lovejoy, 2012), implying that the physical melting and flushing of the ice more strongly govern their presence. Stress due to excessive irradiance increases microalgal susceptibility to chytrid infection in sea ice (Hassett and Gradinger, 2016), and along with salinity stress, these factors may help shape parasitoid succession as host susceptibility varies with environmental factors.

We analyzed 18S ASV time series data for significant correlations between putative parasitoids and potential hosts, assuming correlations could be positive or negative depending on the infection phase captured. It is important to note that correlation does not infer causality and experimental validation is required to confirm real parasite-host associations. Dinoflagellates showed far fewer correlations with putative parasitoids than diatoms, suggesting they are less likely hosts (Fig. 4). The “kill-the-winner” hypothesis postulates that the most abundant taxa experience greater infection rates from viruses or parasitoids, particularly when pathogens are diverse or have a broad host range (Abonyi et al., 2024; Thingstad, 2000). *Navicula* and *Nitzschia* were both highly abundant and significantly correlated with parasitoids, and based on this hypothesis, we would expect more parasitoid associations with these genera. Simpson Diversity was not significantly related to parasitoid abundance, which may reflect uncertainty surrounding the assignment of ASVs as parasitoids, temporal lags between infection and diversity responses, and confounding environmental factors (e.g., brine salinity). A targeted study is required to confirm parasitoid behavior and further investigate whether the suppression of dominant strains by parasitoids may promote microalgal coexistence and diversity in sea ice (Abonyi et al., 2024).

Chytridiomycetes were the most abundant parasitoid group during the algal bloom’s growth phase (Fig. 3a), consistent with previous observations (Donk and Ringelberg, 1983; Ibelings et al., 2004). These host-specific fungal parasites infect Arctic microalgae (Hassett and Gradinger, 2016) and exert top-down control, shaping bloom dynamics and succession. The chytrid ASV, Lobulomycetales Order_1, was significantly correlated with a select few diatoms and dinoflagellates, including *Nitzschia* Genus_4 (Fig. 4). However, their dynamics suggest temporal separation rather than infection (Fig. 5). *Nitzschia* Genus_1 & _5, though not significant (p adj. > 0.05), showed strong correlations (rho ≥ 0.5) and dynamics resembling Lotka-Volterra patterns. *Nitzschia* genera (ASV 1 & 5) were negatively correlated with Lobulomycetales Order_1 during the bloom phase, but were overall, positively correlated, due to post-bloom dynamics. It is therefore likely that we are missing true parasitoid-host interactions that were not significantly correlated while also identifying false positives. Similar inconsistencies in known parasitoid-host pairings have been difficult to parse from high-frequency long-term metabarcoding time-series data due to inconsistent dynamics. We suggest that *Nitzschia* spp. were primary chytrid hosts during our timeseries due to their contrasting peaks and troughs throughout the bloom. Although lifecycle data for marine chytrids are limited, the amphibian pathogen *Batrachochytrium dendrobatidis*, completes its lifecycle in ∼4-5 days (Berger et al., 2005). In our time series, periodic chytrid peaks, and *Nitzschia* genera 1 & 5 dips (Figs. 3 & 5), offset by ∼2 days for genus 5, suggest synchronized infection and zoospore release every ∼7 days. We propose that chytrid peaks reflect sporangia attached to algal hosts retained during sampling, while dips reflect zoospore release and potential loss through brine drainage or dispersal into underlying water. These waves of infection support a cyclic interaction with *Nitzschia* spp. (Fig. 5), though microscopy, isolation, and co-culture are needed to confirm this lifecycle and host range.

Few correlations were found within the Labyrinthulomycetes (Fig. 4); though some thraustochytrids can parasitize diatoms (Hassett, 2020) their typical role is organic matter decomposition (Hassett and Gradinger, 2018). Oomycetes increased post-bloom, along with low levels of Syndiniales (Fig. 3d,e), both of which have a broad host range across Kingdoms (Käse et al., 2021). The order Syndiniales may be more important in underlying waters (Jacquemot et al., 2022; Kellogg et al., 2019; Marquardt et al., 2016; Terrado et al., 2009). Most oomycete ASVs (>85%) were unclassifiable beyond class level, as previously reported in Hassett et al. (2019). Their post-bloom increase may reflect saprotrophic breakdown of ice-retained carbon. However, a negative correlation with a *Cymbellales* ASV, supported by known *Lagena-Cymbella* interaction (Thines and Buaya, 2022), and sequence homology to *Diatomophthora drebesii*, a diatom parasitoid, suggest parasitic roles are probable. Given their relative abundance, Oomycetes may play a key role in Arctic biogeochemical as both degraders and parasitoids.

Cryomonadida (Thecofilosea) featured prominently in our parasitoid correlation matrix (Fig. 4), with some ASVs showing broad host associations among diatoms. *Cryothecomonas* and *Protaspa*, display two trophic modes: heterotrophy of small protists (Thaler and Lovejoy, 2012), and by internal parasitization of large diatoms, with endoparasitism currently restricted to the genus *Cryothecomonas* (Drebes et al., 1996; Kühn et al., 2000). Correlated diatom hosts (e.g.: *Nitzschi*a and *Navicula* spp.) are typically large morphotypes within sea ice, consistent with *Cryothecomonas* hosts. Although *C. aestivalis* is a known diatom parasitoid (Drebes et al., 1996), Arctic *Cryothecomonas* spp. have only been observed as free-living grazers (Thaler and Lovejoy, 2012). In 2022, we observed an undescribed *Cryothecomonas* sp. actively parasitizing *Nitzschia arctica*, consuming the plastids and protoplasm (Fig. 6). *Cryothecomonas* was identified according to Thomsen et al. (1991) and Schnepf & Kühn (2000), i.e., flagellates possess two homodynamic flagella inserted apically, asexual reproduction by binary fission, flagellates were entirely within the host frustule, and the presence of brown digestive vacuoles derived from plastids (Fig. 6). Information regarding the differentiation between the closely related *Protaspa* and *Cryothecomonas* is limited, but current knowledge distinguishes the two based on their feeding behavior, i.e., *Protaspa* attaches to the host diatom, where it remains externally, and penetrates the frustule with a pseudopodia used for feeding (Chantangsi and Leander, 2010; Hoppenrath and Leander, 2006). Endobiotic Oomycetes overtake the diatom protoplasm with a swollen thallus which later develops into a sporangium that releases small biflagellate zoospores (Garvetto et al., 2018); all stages are readily distinguished from *Cryothecomonas*.

Our correlation data suggests that several genera beyond *Nitzschia* and *Navicula* may also be hosts for cryomonads, though *C. aestivalis* is known to be host specific (Drebes et al., 1996; Schnepf and Kühn, 2000) and our observations indicate *N. arctica* was the primary host in 2022 with ∼5% of cells found to be infected*. Protaspa*-host correlations were similar to *Cryothecomonas* suggesting potential host overlap or similar environmental response. As Cryomonadida are abundant in polar waters (Thaler and Lovejoy, 2015; Thomsen et al., 1991), their trophic role in sea ice warrants reevaluation in light of observed parasitization, particularly their influence on diatom dynamics and succession.

We found compelling evidence that the wide diversity of parasitoids observed in this study alter microalgal community composition, particularly of the most abundant sea-ice diatoms. Previous studies have found that, depending on the host species and the parasitoid size classes, carbon flux may be shunted to, or from, grazers (Rasconi et al., 2014). For example, cryomonads and chytrids may redirect carbon from large diatoms—typically too big for grazers like *Calanus glacialis*—into smaller, grazer-accessible forms such as flagellated cells, zoospores, or sporangia (Cleary et al., 2017; Frenken et al., 2016; Rasconi et al., 2014). Parasitoid infections in sea-ice may be prolific (pers. obs.) due to the dense microalgal concentrations in the confined brine channels. Models and experimental investigations indicate enhanced contact rates between viruses and hosts as brine volume decreases with lower temperatures (Wells and Deming, 2006) while meiofaunal grazers become limited in their ability to access brine channels with a diameter less than 200 µm (Krembs et al., 2000). The smaller size of parasitoids, or of certain life stages, are less likely to be limited by narrowing of brine channels, increasing their potential reach and influence within brine channels. Epidemics of parasitoids or viruses may suppress microalgal blooms leading to patchiness, and shift microalgal composition while increasing diversity (Donk and Ringelberg, 1983; Frenken et al., 2016). We emphasize that further research into top-down controls of sea-ice algae and prokaryotes by interactions with parasitoids and viruses is required to better understand community composition structuring and succession.

## 5. Conclusions

As Arctic warming continues, monitoring the timing and magnitude of spring ice-algal blooms is crucial for understanding the ecosystem response to environmental change. High-frequency sampling of landfast sea ice near Utqiaġvik, Alaska in 2021 captured the full bloom cycle, revealing a much larger magnitude relative to prior observations, and highlighting the importance of nitrate limitation in constraining biomass accumulation, and physical forcing (melt onset) on terminating the bloom. Parasitoids emerged as major contributors to the eukaryotic microbial community. We report many potential associations of diatoms with *Cryothecomonas,* including the first observation of parasitic behavior in sea ice, and suggest chytrid-linked declines in *Nitzschia*. Understanding the complex dynamics of parasitoids within the sea-ice ecosystem is crucial to elucidate their role in shaping microbial communities, nutrient cycles, and broader ecological interactions. These findings support the potential role of parasitoids in regulating dominant microalgae through “kill-the-winner” dynamics (Abonyi et al., 2024; Thingstad, 2000), promoting coexistence and microbial diversity by freeing space within the constrained brine channel system for other taxa. As a result, parasitoids may enhance resilience through increased diversity at the base of the sea-ice food web – a critical component of the rapidly changing Pacific Arctic ecosystem.

## Acknowledgements

The authors thank the Ukpeaġvik Iñupiat Corporation Science (UICS) for their logistical support in Utqiaġvik during field work. We thank Hajo Eicken and Rob Rember who were instrumental to securing funding and planning the field campaign, Anika Pinzer for assistance with map making, and Drs. Rebecca Duncan and Jozef Wiktor for their expertise identifying diatoms. This project was supported by research grants funded through the National Science Foundation (NSF) OPP-1735862 (AAI) and OCE-1937715 (GMH) and the Coastal Marine Institute M20AC10007 (GMH), a joint institute between the Bureau of Ocean Energy Management (BOEM) and the University of Alaska Fairbanks. Kenneth Dumack was supported by the German Research Foundation (DFG) with the grant number 555596351. Micrographs were captured at the UAF Molecular Imaging Facility, National Institute of General Medical Sciences (NIGMS) P20GM130443, and sequencing conducted at the UAF Genomic’s Core Lab, NIGMS P20GM103395. We would like to thank our three peer reviewers who volunteered their time and expertise to improve this manuscript.

## Notes

**Conflict of Interest:** The authors declare no conflict of interest.

https://arcticdata.io/catalog/view/doi:10.18739/A21J9793S

https://www.ncbi.nlm.nih.gov/bioproject/?term=PRJNA1108783

https://github.com/kbdilliplaine/2021-Utqiagvik-time-series.git

### Competing Interest Statement

The authors have declared no competing interest.

### Summary of Updates

Considerable revisions were made to main body text and the supplementary materials. Formatting of figures and tables also occurred.

