## Appendix 1 for "Sea-ice microbial community succession and the potential role of parasitoids in the maintenance of diversity during the spring bloom"

Supplemental Methods

*Study site and overview*

The ice formed in situ (refrozen lead) which provided a smooth ice surface with homogenous thickness that is less typical than the rubble ice and small flat pans that characterize the landfast ice in this region. This location was selected due to its distance from shore which minimized dust deposition (~1.5 km) and proximity to a historic sea ice mass balance station.

***Sea ice sampling***

During each sampling event, mean snow depth was measured (n=5 within 1 m^2^) and the temperatures of air, snow surface, snow/ice interface and ice were quantified using a Fisher Scientific Traceable thermometer (± 0.05 °C). Biological samples, i.e., chl *a* and DNA, were collected from a single 10-cm section of bottom ice which was sectioned using an ethanol sterilized handsaw immediately upon core extraction. Samples from DOYs 120 and 139 differed from the 10 cm length and were 12 and 5-cm section lengths, respectively. Ice thickness of each core hole was recorded using a thickness gauge and reported as an average of all collected cores for each sample day (Fig. S2c; n=2-9).

***Physical, chemical and biological measures***

Ice temperature was measured at 5-cm intervals (Oggier et al., 2020). Temperature and salinity of the seawater were measured through a core hole approximately 1 m below the ice. We defined ice growth/melt periods by whether the temperature of the ice measured 2.5 cm from the bottom was above or below the freezing point of seawater. The freezing point of seawater was calculated as -1.8 °C by assuming a salinity of 32.4, which was the average seawater salinity of the first 25 days before freshening due to sea ice melt was observed (Kelley and Richards, 2022). A separate nutrient and salinity core was sectioned into 5-cm sections immediately upon retrieval, working inward from both ends (Oggier et al., 2020). Macronutrient samples were extracted by melting each section in sealed acid-cleaned jars at room temperature in the dark and filtered within 24 hours. Seawater was collected using a plastic hand-operated pump through a hole in the ice and stored in the dark at 4 °C for no more than 24 hours prior to filtration. Nutrient samples were processed for nitrate + nitrite, nitrite, phosphate and silicate. Remaining sample was used to measure the bulk salinity using a YSI 30 handheld conductivity meter. Density was measured from a separate core. The values for nutrients, temperature, salinity, and density from the bottommost two 5-cm sections of cores were averaged to provide a single value representative of a 10-cm section to match the biological measures. Linear regressions were used to determine the correlation of bulk nutrient concentrations with areal chl *a* (Fig. S5).

Ice temperatures were measured immediately upon retrieval from one or more cores at 5-cm intervals starting from each end of the core with a 2.5-cm offset from the ends (Oggier et al., 2020). Brine volume and salinity were calculated according to the equations of Cox and Weeks (1983). Five of the field collection days did not have accompanying density data, and no trend was seen across the timeseries, so the average from the entire field campaign was used (908 kg m^-3^; Day of Year (DOY) 123, 126, 134, 139 & 145). Nutrient cores were processed proceeding from each end so that a variable length core section was always confined to the interior of the ice. Nutrient samples from melted ice and seawater were filtered through 0.45 µm cellulose acetate filters into 20-mL acid-cleaned HDPE containers and frozen at -20 °C until analysis. Nutrient concentrations (nitrate + nitrite, nitrite, phosphate, and silicate) were determined by colorimetry using a Seal Analytical continuous-flow QuAAtro39 AutoAnalyzer following Becker et al. (2020). Prior to analysis, frozen samples were heated to 50 °C for 30 minutes, then allowed to come to room temperature. Due to the variable salinity of the samples, standards were salinity-adjusted with Milli-Q water to match the salinity of the samples being analyzed. Silicate was determined according to Armstrong (1967). Phosphate was analyzed following modifications of Murphy and Riley (1962). Nitrate was determined by reducing nitrate to nitrite following Armstrong et al., (1967). Certified Reference Materials (KANSO-CRM) were included with each analytical run. Nutrients from ice are presented as brine concentrations (Fig. S4b). Brine concentrations were calculated using the averages for the bottom 10 cm of ice (usually two segments for nutrients and density). The melted volume was adjusted by multiplication of the measured ice density as a fraction of liquid water density and scaled to the brine volume fraction.

***Chlorophyll a and Phaeophytin analysis***

Algal pigments were extracted from filters with 7 ml of 90% acetone at -20 °C for 24 hours; samples were warmed to room temperature for 1 hour in the dark before fluorometric measurement using a Turner 10-AU fluorometer, before and after acidification, to determine pigment concentrations (chlorophyll *a* and phaeophytin; Arar and Collins 1997). Core samples with section lengths different from the bottom 10-cm section (DOYs 120 and 139) were excluded from brine concentration calculations. A Weibull distribution curve was fit to areal chl *a* concentration. The bloom peak and end dates, were calculated from the chl *a* models and defined as the global maximum (peak) and when concentration decreased below 1 mg m^-2^ (end; Leu et al., 2015). We categorized our samples within two phases of the sympagic algal bloom, i.e., “bloom” chl *a* >1 mg m^-2^ and “post-bloom” <1 mg m^-2^. Chl *a* brine concentration was calculated in the same manner as the nutrients (Section 1.2), after subtraction of the added filtered seawater volume. A lognormal distribution curve was fit to brine chl *a* concentration (Fig. S5).

***Irradiance measurements and estimations***

To account for the effect that time of day and cloud cover had on *in situ* measurements, we calculated the 4π:2π ratio, referred to in this paper as “pseudotransmissivity,” represented as a percent, when simultaneous measures were available. Long-term incident PAR data were also recorded at the nearby National Ecological Observatory Network, Utqiaġvik, Alaska (BARR), station as an average at 30-min intervals (Fig. S2a; NEON, 2023). Percent pseudotransmissivity was linearly interpolated between sampling days to fill gaps in data and multiplied by incident NEON irradiance to calculate higher-frequency irradiance at the ice-water interface (Fig. S2e). We also modelled the irradiance at the ice-water interface using the Lambert Beer equation, with and without chl *a* (Fig. S2g,h):

$$E_{z}={E_{0}}^{{(-K}_{s}*z_{s}){+(-K}_{i}*z_{i})+{(-K}_{chl}*[chl a])}$$

where E_z_ is the irradiance at the ice-water interface, E_0­_ is the incident irradiance measured at NEON, and the diffuse attenuation coefficients of snow (K_s_= 11), ice (Ki, 3.6), and areal chl *a* concentration (K_chl_, 0.06) were derived from McDonald et al. (2015), z is the thickness of snow or ice and [chl *a*] is the areal chl *a* (mg m^2^). As in Dilliplaine and Hennon (2023), K_i_ was adjusted to 2.6, within the reported confidence interval, to allow for upwelling irradiance as measured *in situ*.

***Amplicon generation, sequencing, and bioinformatic processing***Sterivex filters were sliced and removed from their housing using ethanol cleaned scalpel and forceps prior to DNA extraction following the NucleoMag DNA/RNA water kit according to filters sized 47 mm (Macherey Nagel, Düren, GE). 16S rRNA gene amplicons were produced using the revised Earth Microbiome Project primers 515F (5’-GTGYCAGCMGCCGCGGTAA-3’) and 806RB (5’-GGACTACNVGGGTWTCTAAT-3’) (Caporaso et al. 2012; Apprill et al. 2015; Parada et al. 2016). 18S rRNA gene amplicons were generated using the primers TAReuk454FWD1 (5′-CCAGCASCYGCGGTAATTCC-3′) and TAReukREV3 (5′-ACTTTCGTTCTTGATYRA-3′) according to reaction recipe and thermal cycling conditions in Appendix 1 (Stoeck et al., 2010). Polymerase chain reaction recipes were performed in 25 ul reactions with the following composition using Kapa BIOSYSYSTEMS HiFi Hotstart Polymerase (kk2502): 16.25 µL dH_2_O, 5 µL 5X Buffer, 0.75 µL 10 mM dNTPs, 0.75 µL 10 mM of forward and reverse primers, 0.5 µL Taq polymerase, and 1 µL of gDNA. Thermocycling conditions included a 1 minute denaturation at 98 °C, followed by 25 cycles of 98 °C for 20 s, 55 °C for 20 s, and 72 °C for 30 s, followed by a 5 minute final elongation at 72 °C.

Sample libraries were initially demultiplexed and outer barcodes were removed as part of the standard Illumina post-processing. Sequenced samples were demultiplexed, and internal indices removed, using the Mr. Demuxy package (Glenn et al., 2019). First, primer regions were removed using Cutadapt (Martin, 2011). Sequences were denoised and dereplicated using DADA2, including quality control, chimera checking and paired-end joining in R (Callahan et al., 2016). Base qualities were visually assessed, and sequences trimmed, when quality score of the 25^th^ percentile reached 33. Denoising was conducted on “pseudo”-pooled samples for increased sensitivity. Chimeras were identified and removed from samples individually with a “—p-min-fold-parent-over-abundance” value of 2. The R package ‘microViz’ was used to standardize taxonomic naming when corrections were required with the “tax_fix()” command (Barnett et al., 2021). Non-target amplicons such as 16S sequences belonging to eukaryota, chloroplasts, mitochondria, or unassigned, and 18S sequences belonging to bacteria, metazoa, unassigned, or plastid DNA, were removed. Phyloseq was used for data manipulation and rarefaction prior to analyses; either rarified or unrarefied data were used dependent on the analysis. Prokaryotic and eukaryotic sequence data were rarefied to 10,497 and 15,684 reads per sample, respectively. Unrarefied data were converted to within sample relative abundance.

Following Walker et al. (2023), community structuring was investigated using the “ward.D2” method for hierarchical clustering was used to identify unique clusters (Murtagh and Legendre, 2014). The ‘vegan’ package was used to investigate cluster validity, within-cluster dispersion, and significant differences in composition between clusters using a silhouette test, dispersion test “betadisper,” and PERMANOVA “adonis2”, respectively (Oksanen et al., 2008).

All *Tryblionella* ASVs were reassigned to the genus *Nitzschia* and *Amphiprora* ASVs to the genus *Entomoneis*. All other changes were simple decreases in taxonomic resolution, e.g., *Attheya septentrionalis* to *Attheya* sp, as our small fragment of the 18S gene is not sufficient to identify organisms to the species level in many cases.

***Microscopy***

Samples obtained from the bottom 5-cm of ice, collected on April 25, 2022, were melted according to the biological methods in section 2.1 and transported to the University of Alaska Fairbanks in a cooler with ice in the dark. Live material was viewed using differential interference contrast microscopy (Nikon Eclipse 80i with View 4k Color camera) at 400 – 1000x magnification (Fig. 6). Healthy and *Cryothecomonas-*infected *Nitzschia arctica* cells were enumerated in formaldehyde-fixed samples collected at the same site on April 26, 2022 using a Sedgewick-Rafter chamber at 400x magnification on a Nikon TE2000U inverted microscope. This material was collected as triplicate subsamples of the bottom 2-cm of twelve 20-cm inner diameter cores pooled together and concentrated on a 4 µm mesh sieve. A minimum of 500 *N. arctica* cells were counted per subsample.

Supplemental Figures

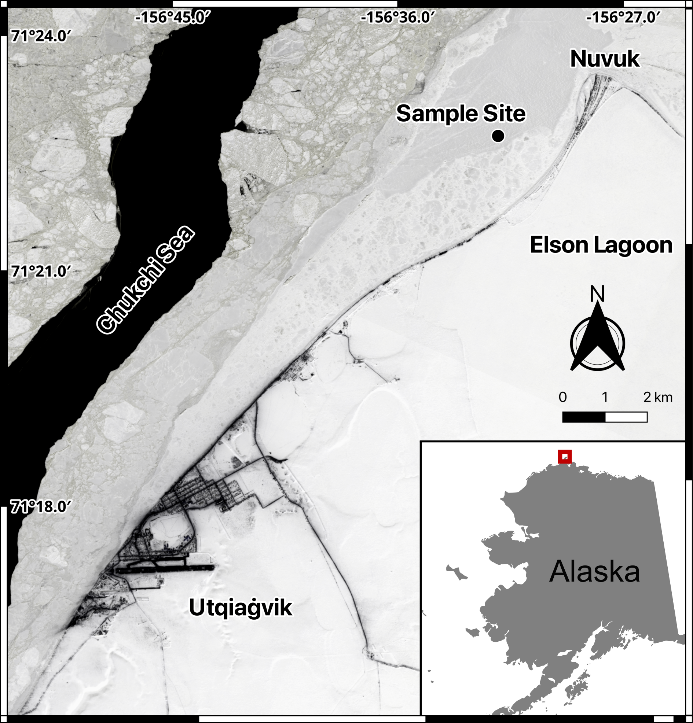

**Supplementary Figure S1.  Location of the sample site near Utqiaġvik, Alaska**. Red square on the inset map shows the location of Utqiaġvik, AK. The 4-band reflectance Planet image was taken on April 25^th^, 2021.

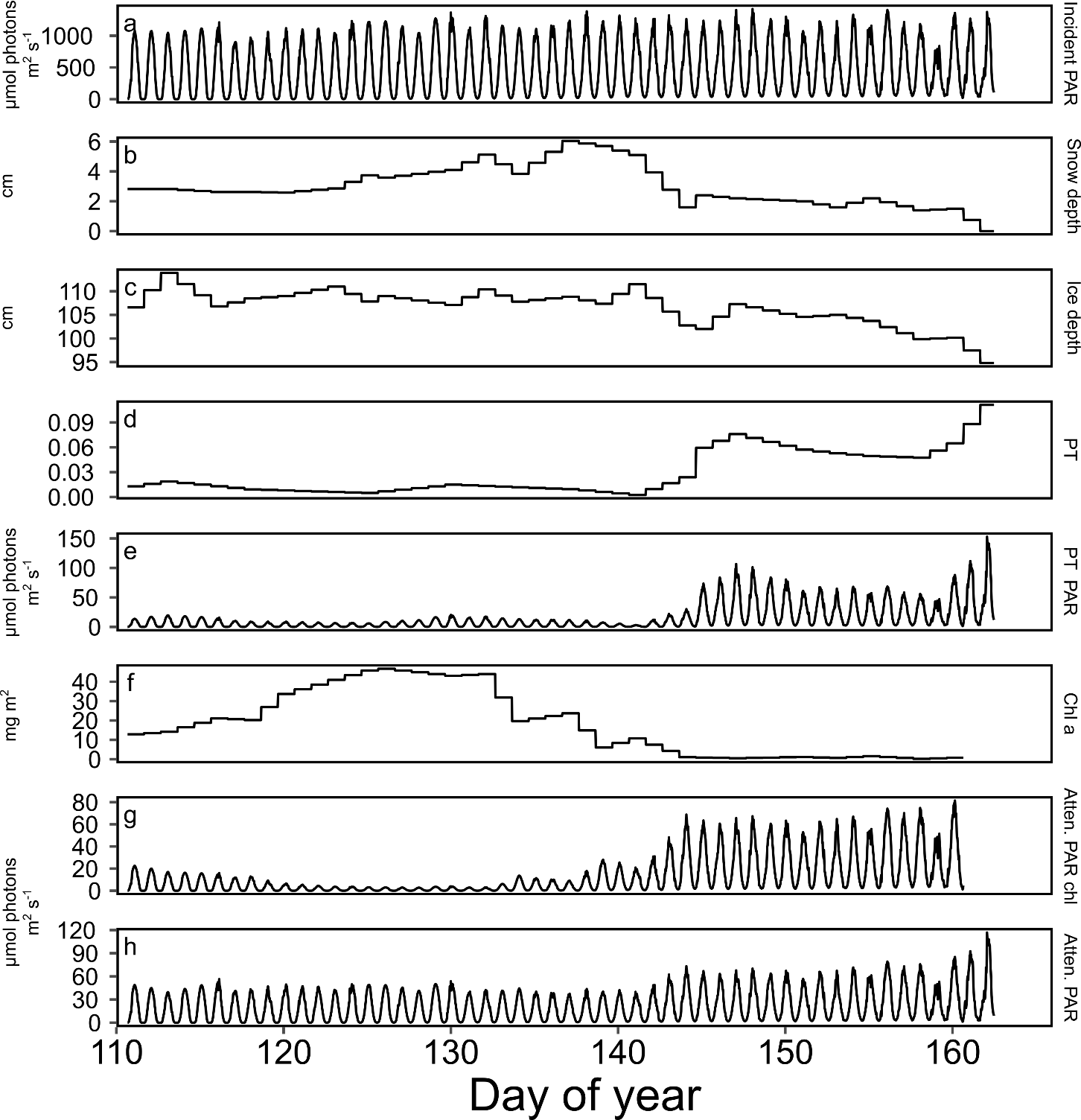

**Supplementary Figure S2. Light, attenuating factors, and calculated attenuation coefficients during the sampling campaign.** Incident photosynthetically active radiation (PAR; 400-700 nm) measured at the National Ecological Observatory Network in Utqiaġvik, AK (a), snow depth with linear interpretations between measurements (b), ice thickness with linear interpretations between measurements (c), “pseudotransmissivity” as the ratio of *in situ* incident irradiance and under-ice irradiance (PT; see methods 2.3 for description) with linear interpretations between measurements (d), pseudotransmissivity calculated PAR (PT PAR) as the multiplication of the PT ratio by NEON measured incident irradiance (e), chlorophyll *a* with linear interpretations between measurements (f), calculated under-ice irradiance using the attenuation of snow and ice, with (g) and without (h) chlorophyll *a*.

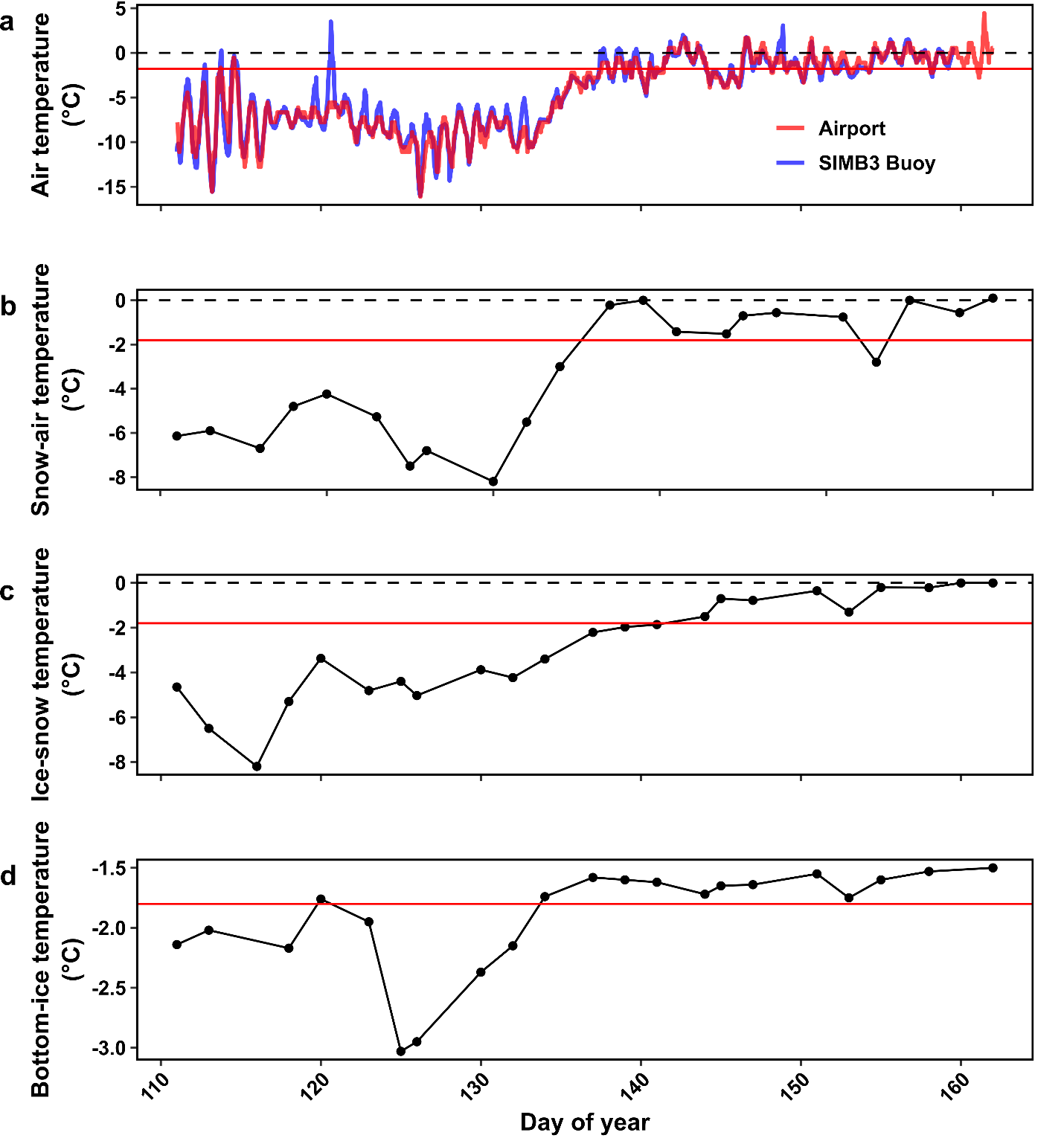

**Supplementary Figure S3. Ambient, interface, and bottom-ice temperatures** **during the sampling campaign.** Air temperature measured at continuous recording stations in the town of Utqiaġvik (Airport) and on the ice at a nearby sea-ice mass balance station (SIMB3) (a), temperature measured in the field at the snow-air interface, (b) the ice-snow interface (c), and measured 2.5-cms from the ice bottom (d). Dashed black line is 0 °C isotherm and the red solid line is the -1.8 °C isotherm, representing the melting points of fresh and saltwater ice, respectively.

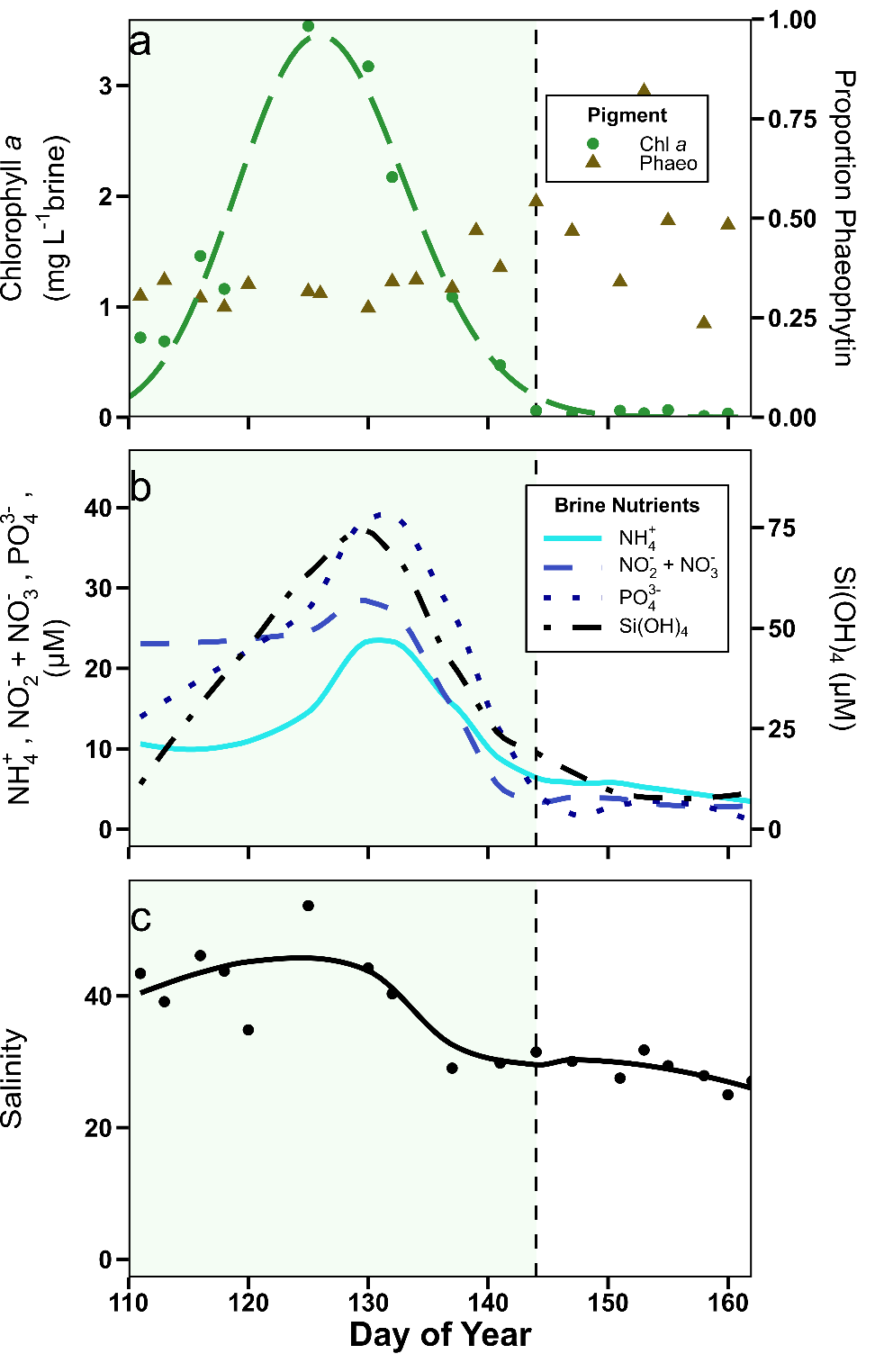

**Supplementary Figure S4. Temporal evolution of calculated brine concentrations from the bottom 10-cm of ice.** Chlorophyll *a* and phaeophytin proportion (a), ammonium (NH_4_^+^), nitrite + nitrate (NO_2_^-^ + NO_3_^-^), phosphate (PO_4_^3-^_­_) and silicate (Si(OH)_4_; b), and salinity (c). Green shading indicates the ice-algal bloom and the vertical dashed line indicates the bloom termination (<1 mg chlorophyll *a* m^-2^).

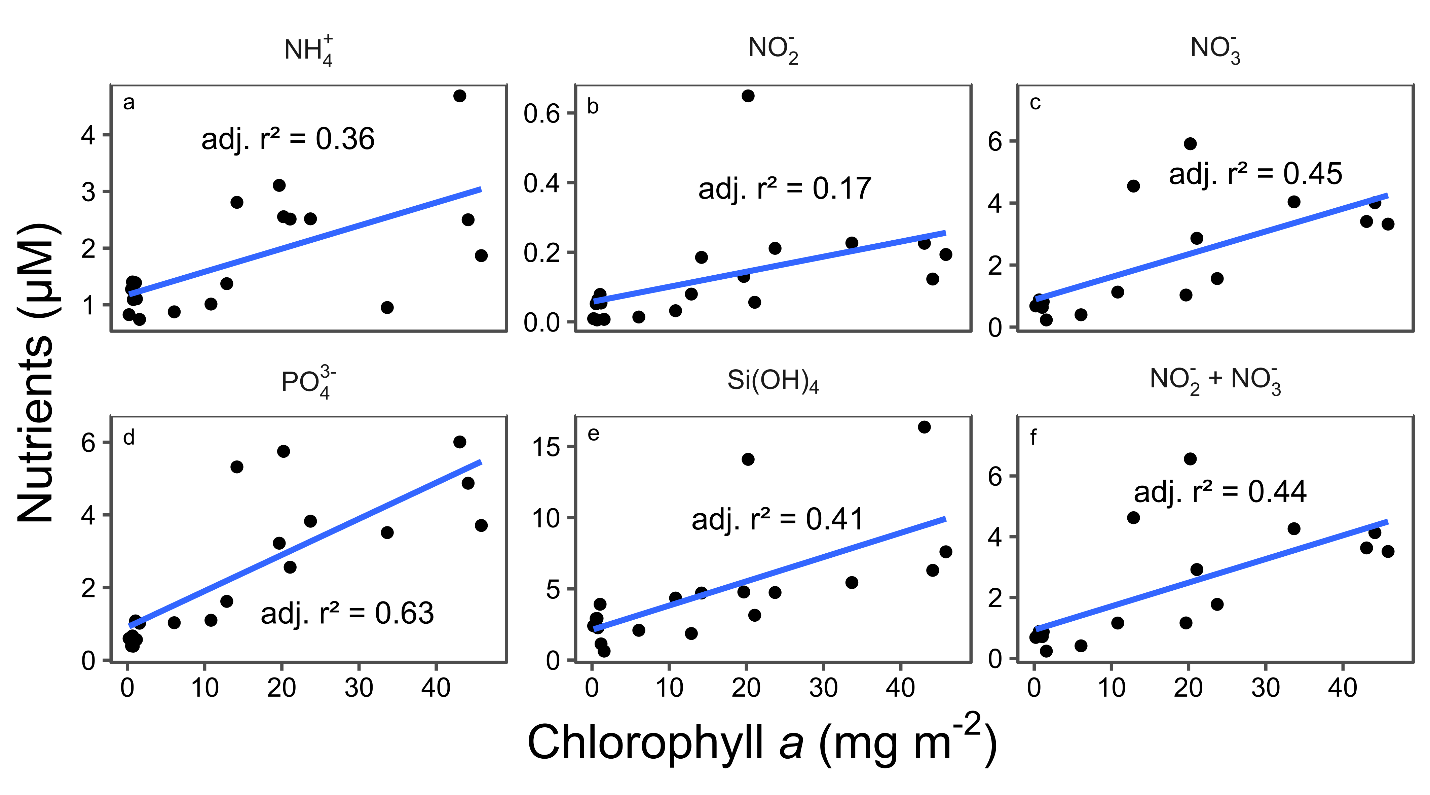

**Supplementary Figure S5. Linear regressions of bulk nutrient concentrations vs. areal chlorophyll *a* concentration.** Ammonia (a; NH_4_^+^), Nitrite (b; NO_2_^-^), Nitrate (c; NO_3_^-^), Phosphate (d; PO_4_^3-^), Silicate (e; Si(OH)_4)_ and total nitrogen (f; NO_2_^-^ + NO_3_^-^).

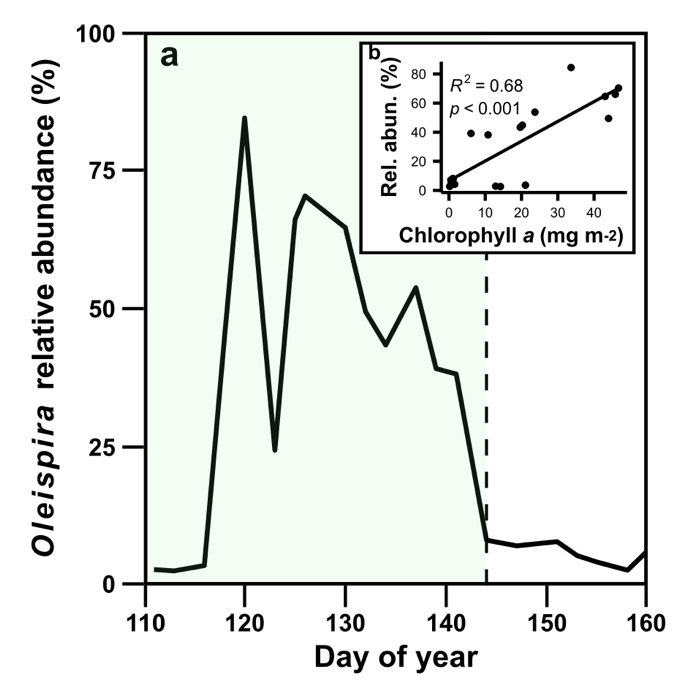

**Supplementary Figure S6. Relative abundance of the genus *Oleispira* over time (a) and its correlation to chlorophyll *a* (b).** Green shading indicates the ice-algal bloom and the vertical dashed line indicates the bloom termination (<1 mg chlorophyll *a* m^-2^).

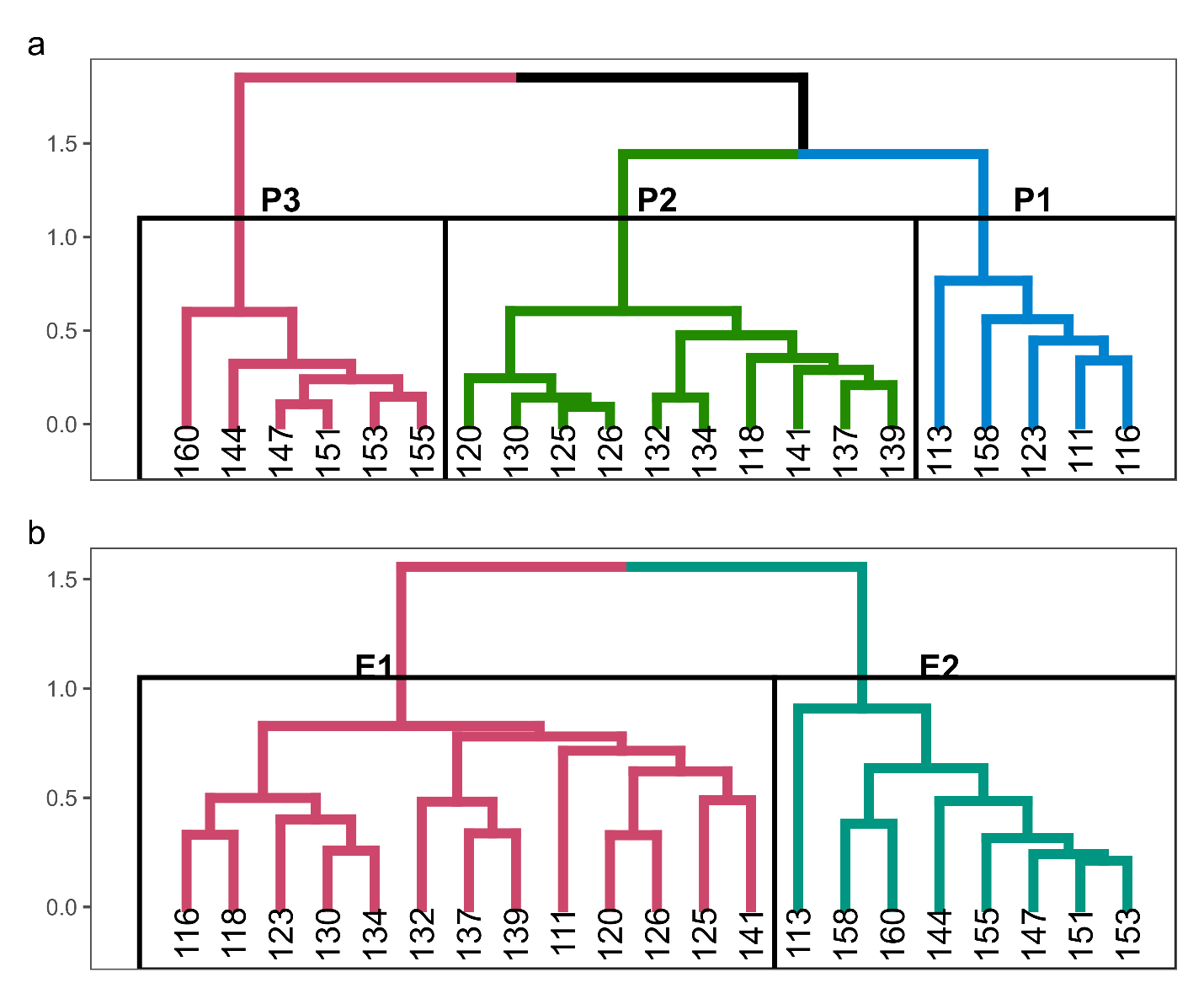

**Supplementary Figure S7.** **Dendrogram of clustering results based on 16S rRNA (a) and 18S rRNA (b) community composition.** Each node tip is a sample day.

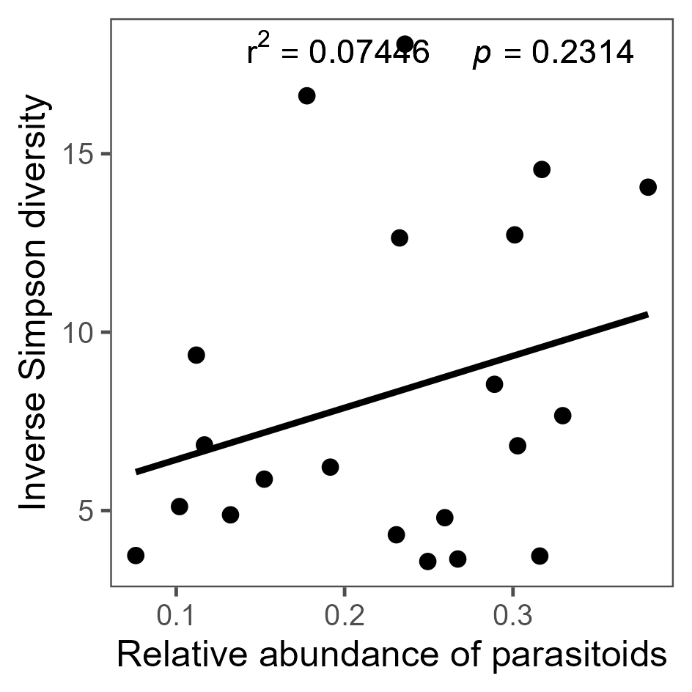

**Supplementary Figure S8. Linear regression of microalgal Inverse Simpson diversity with the total sum of parasitoid class relative abundance.** Microalgal taxa were limited to the Bacillariophyceae, Mediophyceae, Coscinodiscophyceae, and Dinophyceae and the summed relative abundance of putative parasitoids were limited to the classes Chytridiomycetes, Thecofilosea (Cercozoa), Labyrinthulomycetes (Bigyra), Oomycetes and Syndiniales (Dinoflagellata).

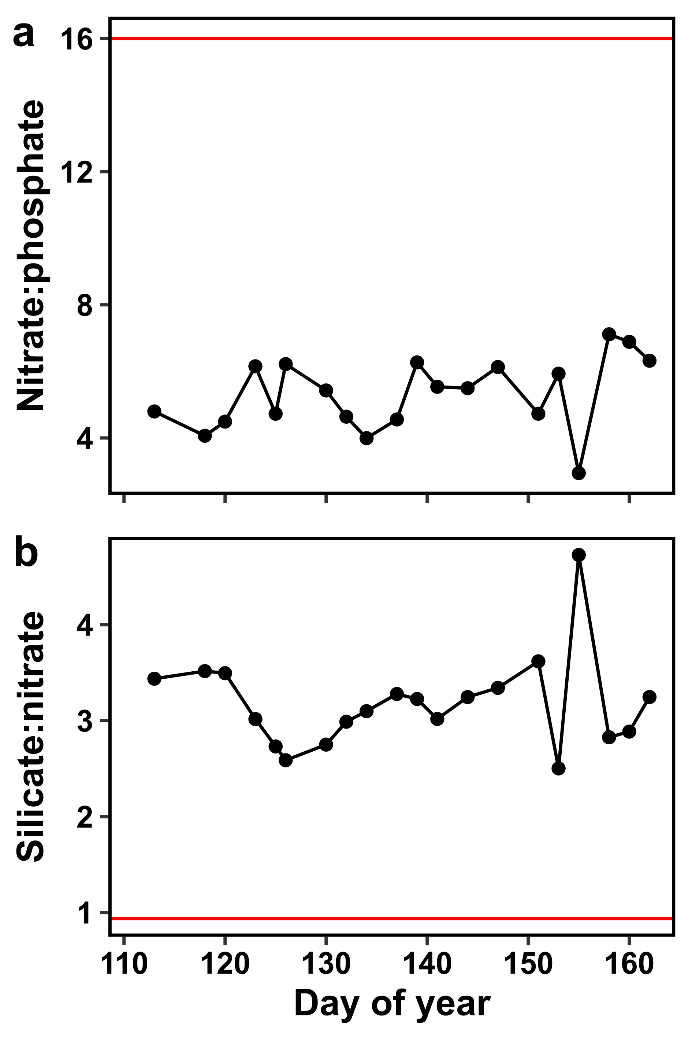

**Supplementary Figure S9. Seawater nutrient ratios of nitrate:phosphate (a) and Silicate:nitrate (b)** with idealized Redfield ratios as the horizontal redlines of 16:1 (N:P) and 16:15 (Si:N).
